# Identification and characterization of a G-quadruplex structure in the pre-core promoter region of hepatitis B virus

**DOI:** 10.1101/2021.01.19.427307

**Authors:** Vanessa Meier-Stephenson, Maulik D Badmalia, Tyler Mrozowich, Keith CK Lau, Sarah K Schultz, Darren L Gemmill, Carla Osiowy, Guido van Marle, Carla S Coffin, Trushar R Patel

## Abstract

Worldwide, ∼250 million people are chronically infected with the hepatitis B virus (HBV) and are at increased risk of cirrhosis and hepatocellular carcinoma. The HBV persists as covalently closed circular DNA (cccDNA), which acts as the template for all HBV mRNA transcripts. Nucleos(t)ide analogs do not directly target the HBV cccDNA and cannot eradicate the HBV. We have discovered a unique structural motif, a G-quadruplex in HBV’s pre-core promoter region that is conserved amongst nearly all genotypes, and is central to critical steps in the viral life-cycle including the production of pre-genomic RNA, core and polymerase proteins, and encapsidation. Thus, an increased understanding of the HBV pre-core may lead to the development of novel anti-HBV cccDNA targets. We utilized biophysical methods to characterize the presence of the G-quadruplex, employed assays using a known quadruplex-binding protein (DHX36) to pull-down HBV cccDNA, and compared HBV infection in HepG2 cells transfected with wild-type and mutant HBV plasmids. This study provides insights into the presence of a G-quadruplex in the HBV pre-core promoter region essential for HBV replication. The evaluation of this critical host-protein interaction site in the HBV cccDNA may ultimately facilitate the development of novel anti-HBV therapeutics against the resilient cccDNA template.

## INTRODUCTION

Approximately 250 million people worldwide are chronic hepatitis B virus (HBV) carriers and are at elevated risk of developing cirrhosis, liver failure, and hepatocellular carcinoma (HCC) (1–3). The virus persists, in part, due to the presence of its stable, compact intranuclear minichromosome— the covalently closed circular DNA (cccDNA) (4–7). Current oral therapies against HBV, including nucleotide/side analogues (NA) target the virus during replication and are effective at decreasing viremia, but rarely induce HBV surface antigen loss, i.e., “functional cure” (8–11). Moreover, NAs do not directly target the cccDNA and can lead to viral relapse once treatment is stopped (10–13). Achieving a sterilizing (or virological) HBV cure requires targeting and clearance of the HBV cccDNA episome. There is limited knowledge of the secondary and tertiary structural intricacies of the HBV genome. The discovery of unique HBV cccDNA structural motifs may highlight new areas for therapeutic drug design and intervention.

HBV is a small, compact 3.2 kb, partially double-stranded DNA virus (14). The HBV genome consists of four overlapping open-reading frames for genes surface (S), core (C), polymerase (P), and X, encoding seven viral proteins (14). Transcriptional regulation of these regions is guided by the corresponding promoter regions that enable the binding of various cellular (and viral) proteins to initiate transcription and produce five RNA transcripts, including the pre-C and pre-genomic RNAs (pgRNA) (each approximately 3.5 kb), and the sub-genomic RNAs S1 (2.4 kb), S2 (2.1 kb) and X (0.7 kb) (14). The pre-C / pre-genomic region is one of the most important viral promoter regions. This region initiates transcription of the mRNA template for the production of both the core protein and polymerase and produces the full-length pre-genomic RNA which contains the 5 -hairpin loop or epsilon (ε) (14–16). The epsilon region ′ ε enables binding of the viral polymerase and initiation of encapsidation or packaging of the viral template followed by replication and the creation of viral progeny (14, 15). Overall, HBV is classified into eight main genotypes, A-H, based on a >8% nucleotide variation across the genome (17, 18). The genotypes tend to display different propensities to cause clinical disease and to respond to therapy (19, 20). Despite the number of unique genotypes and viral variants due to the HBV’s error-prone reverse transcriptase, promoter regions tend to be highly conserved amongst all HBV variants / genotypes (21). This is likely owing to their need to interact with key host proteins for successful propagation.

G-quadruplexes are highly stable non-canonical DNA (or RNA) structures formed in guanosine-rich nucleic acid regions, where four guanines form a planar quartet (G-quartet) stabilized by the presence of Hoogsteen-hydrogen bonds (22). Three or more consecutive G-quartets can stack on top of each other to form parallel, anti-parallel, or hybrid G-quadruplex structures (23). The G-quadruplexes are further stabilized by monovalent cations that occupy the central channel between the G-quartets (22). The typical sequence that can produce a quadruplex is represented by the formula, -G_3_-N_1-7_-G_3_-N_1-7_-G_3_-N_1-7_-G_3_-, where groups of three guanosines are separated by one to seven nucleotides, but there are exceptions to this rule (24), such as the specific quadruplex highlighted in the current study. Over 700,000 putative quadruplex sequences have been found throughout the human genome, having a predominance in the telomeric and gene regulatory regions, which has led to the discovery of their roles in key transcription, translation, and genomic regulatory processes (25–31). In the field of cancer research, such findings have facilitated the development of a new class of drugs (*i.e*., G-quadruplex stabilizers), including the recent telomeric quadruplex stabilizers, CX-3545 (Quarfloxin®) and CX-5461, currently in early Phase clinical trials (32–35). Based on advances in oncology, quadruplexes have subsequently been found in several viral genomes, including human immunodeficiency virus-1 (HIV-1) (36), herpes simplex virus (HSV-1) (37, 38), hepatitis C virus (HCV) (39), human papillomavirus (HPV) (40), Epstein-Barr virus (EBV) (41), Kaposi’s sarcoma-related herpesvirus (KSHV) (42), and Zika virus (43); reviewed in (44, 45). More recently, a G-quadruplex has also been found in the promoter region of the HBV’s S-gene (46). The predominance of G-quadruplexes in key regulatory regions is highlighted, suggesting interesting targets for anti-viral therapy development (44, 45).

In the current work, we demonstrate the central, multi-functioning pre-core promoter region of the HBV genome as a G-quadruplex forming region, confirm its presence through biophysical methods, and demonstrate its role in viral replication when compared to non-G-quadruplex-forming mutants *in vitro*. Collectively, this lays the basis for the study of a critical host-protein interaction and the potential development of unique therapeutic targets for HBV cccDNA.

## RESULTS

### The HBV pre-core promoter region contains a highly conserved G-rich sequence

In our prior analysis of the HBV promoter region using the HBV genome database (HBVdB: https://hbvdb.ibcp.fr), we noted a highly G-rich region in the pre-core/core promoter region in all HBV genomes except the HBV G genotype (21). This region would be overlooked if the rigid formula were used when searching for putative quadruplex sequences, G_3-5_ N_1-7_ G_3-5_ N_1-7_ G_3-5_ N_1-7_ G_3-5_, (i.e., where G’s are guanosines present in groups of three to five, and N’s are any nucleotide present in one to seven sequences separating these groups of guanosines), but exceptions exist (47). This region was of significant interest given the fact that it also binds to host Specificity protein 1 (Sp1), a host transcription factor already known to bind G-quadruplexes, further supporting the link with the secondary structure in this region. The analysis of single nucleotide mutation frequencies in this region was computed similarly to our previous study (21), but with a focus on all available basal core promoter sequences in the HBV genome database (9939 sequences, Figure 1A), demonstrating remarkable conservation of the guanosine groups.

**Figure 1.**
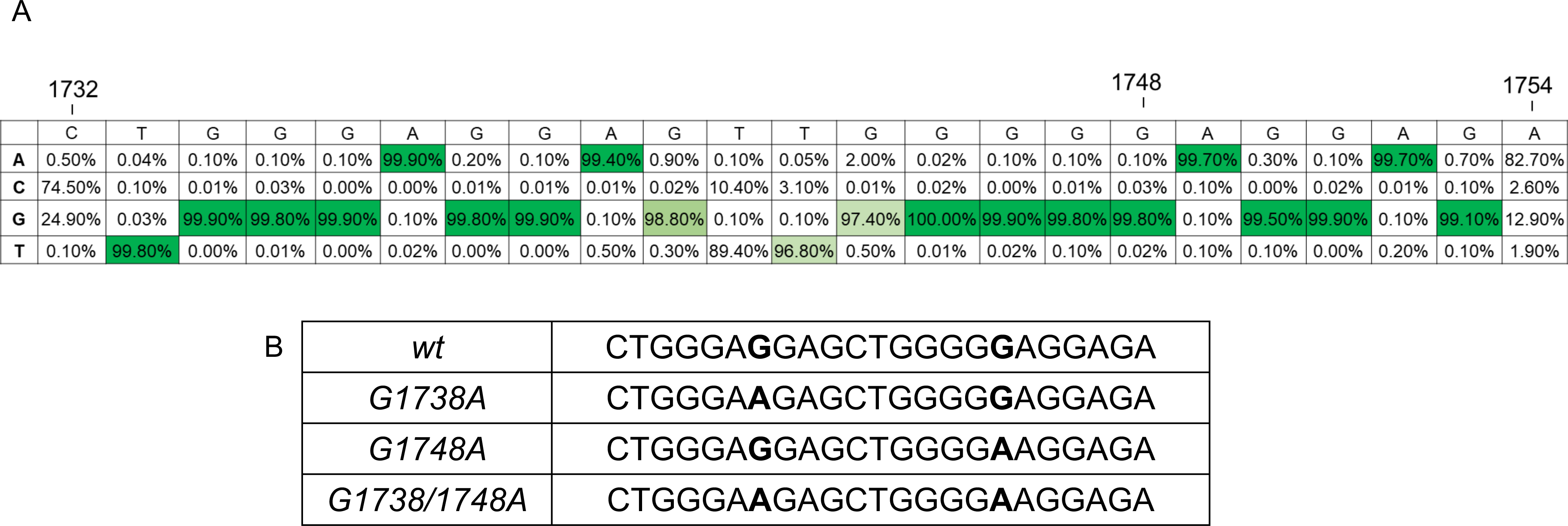
(A) Frequency analysis of the guanosines present in the pre-core promoter region of HBV using the HBV genome database https://hbvdb.ibcp.fr, (accessed April, 2019). HBV Genotype G does not contain this G-rich pre-core region and hence was not included in the analysis. (B) Wild-type (*wt*) and mutants (*G1738A*, *G1748A* and *G1738/1748A*) oligomers used in this study.

### The pre-core/core promoter region of HBV forms a quadruplex structure

To determine the ability of the HBV pre-core/core promoter region to form a quadruplex, we utilized a 23-mer oligomer of the wild-type (*wt*) promoter region for use in multiple biophysical assays (Figure1B). We also designed a mutant (*G1748A*) oligomer based on prior studies showing loss of Sp1 binding with the single-nucleotide mutation of G to A substitution at position 1748 (48). Additional mutants were also included (*i.e.*, *G1738A* and *G1738/1748A*) to determine the nature and necessity of these putative G-quadruplex-disrupting mutations. HBV genome frequency analysis using the HBV database (www.hbvdb.ibcp.fr) showed >99.8% conservation across all genotypes hence supporting the use of these mutants for our proposed studies (Figure 1). Oligomers were solubilized in an appropriate buffer, followed by a heat- cooled step to allow for G-quadruplex formation. Samples were purified using SEC prior to performing all experiments to ensure they were free of aggregation and showed a single peak eluted for subsequent collection and concentration (Figure 2A). First, we performed circular dichroism spectropolarimetry (CD) experiments to investigate whether the wildtype (*wt*) and mutant oligomers form a G-quadruplex structure in solution. We observed a peak in ellipticity upon CD analysis at λ≈263nm and a minimum at λ≈ 242 nm (as detailed in Table S1, Figure 2B), suggesting that the *wt* oligomer adopts a parallel G-quadruplex structure in solution (49–51). Similarly, the *G1738A* mutant showed a comparable quadruplex CD profile. The spectra for the *G1748A* and *G1738/1748A* mutants, however, displayed a less prominent pattern despite the same concentration, consistent with a partially folded quadruplex structure (Figure 2B). We also studied the G quadruplex formation in the presence of Li^+^. We purified G-quadruplexes under identical conditions except that the buffers contained Li^+^ instead of K^+^. The CD spectra were collected for all the samples, and no considerable changes were observed in CD spectra collected for G-quadruplexes in Li^+^ and K^+^, suggesting that the DNA sequence can form G-quadruplex despite possessing two mutations irrespective of the centrally coordinated monovalent cation. Therefore, we decided to collect MST and SAXS data for G quadruplexes in the presence of K^+^ only.

**Figure 2.**
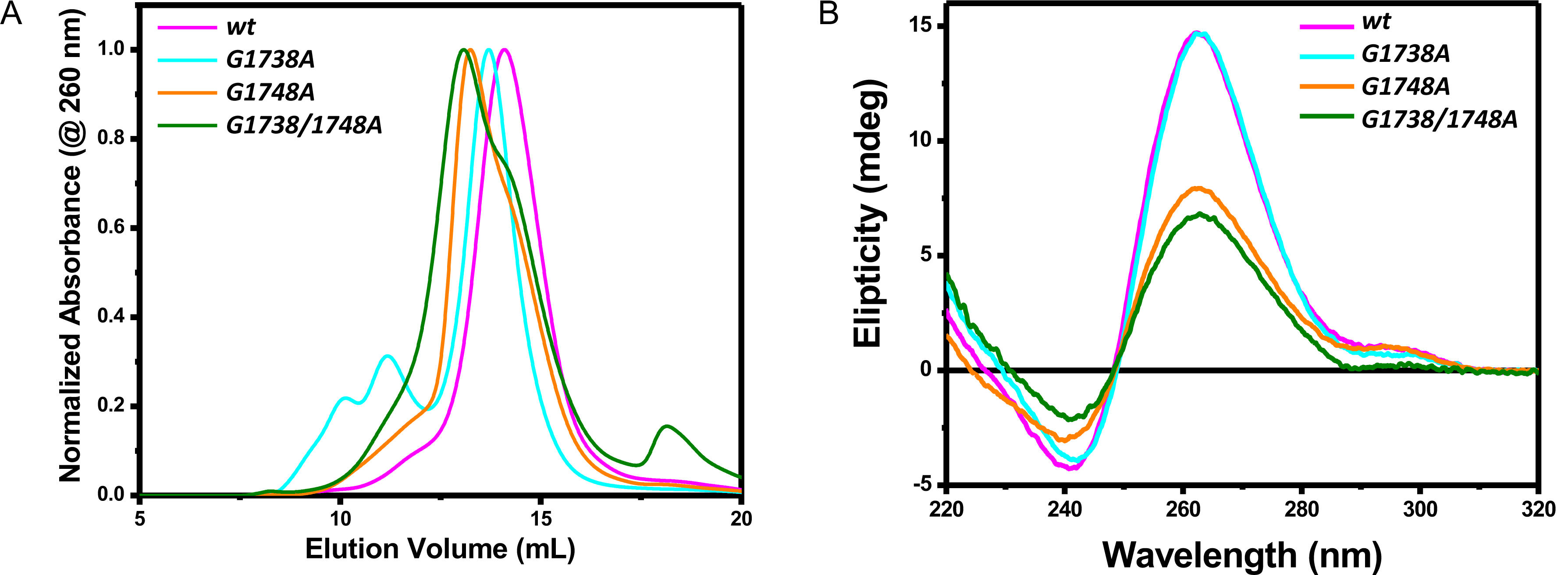
Purification and primary biophysical studies of the pre-core promoter *wt*, *G1738A*, *G1748A* and *G1738/1748A* mutant oligomers. (A) The change in size exclusion chromatography elution profile suggests change in size of G4s (B) Circular dichroism spectroscopy studies of the pre-core promoter oligomers. The presence of a negative peak at ∼242 nm and a positive peak at ∼263 nm indicates that the G4s are present at a parallel quadruplex in the solution, the respective values are tabulated in Table S1. The plots represent the average of the three independent runs for each.

**Table 1:** Analysis of small-angle X-ray scattering data for HBV wild-type (*wt*) and mutant (*mut*) core promoter oligomer samples

|  | *wt* | *G1738A* | *G1748A* | *G1738/1748A* |
| --- | --- | --- | --- | --- |
| $M_w$ (kDa) | 7.30 | 7.28 | 7.28 | 7.26 |
| $I(0)^\#$ | $0.0035 \pm 1.1 \times 10^{-5}$ | $0.0041 \pm 1.3 \times 10^{-5}$ | $0.0031 \pm 1.6 \times 10^{-5}$ | $0.0037 \pm 2.1 \times 10^{-5}$ |
| $q.R_g$ range | 0.24 – 1.30 | 0.23 – 1.30 | 0.23 – 1.30 | 0.23 – 1.29 |
| $R_g$ (Å)<sup>#</sup> | $18.37 \pm 0.11$ | $19.10 \pm 0.11$ | $20.10 \pm 0.19$ | $20.34 \pm 0.22$ |
| $I(0)^{\#\Delta}$ | $0.0034 \pm 0.7 \times 10^{-5}$ | $0.0039 \pm 1.0 \times 10^{-5}$ | $0.0030 \pm 1.2 \times 10^{-5}$ | $0.0036 \pm 1.7 \times 10^{-5}$ |
| $R_g$ (Å)<sup>Δ</sup> | $17.18 \pm 0.03$ | $18.17 \pm 0.04$ | $19.77 \pm 0.09$ | $20.43 \pm 0.10$ |
| $D_{max}$ (Å)<sup>Δ</sup> | 46.85 | 53.14 | 60 | 60.44 |
| $\chi^2$* | ~1.80 | ~1.50 | ~1.50 | ~1.20 |
| NSD* | $0.55 \pm 0.01$ | $0.58 \pm 0.03$ | $0.58 \pm 0.01$ | $0.59 \pm 0.01$ |

The molecular weight ( $M_w$ ) values were calculated using nucleotide sequences.
<sup>#</sup> - obtained from Guinier analysis (89).
<sup>Δ</sup> - determined using $P(r)$ analysis using the GNOM program (55).
\* - values derived from DAMMIN (90) and DAMAVER (84) analysis.
The error calculation for each SAXS data analysis step is described by individual packages implemented in ATSAS suite (55, 84, 89, 90).

To visualize and verify the G-quadruplex formation, we employed the HPLC-SAXS (52, 53). This enabled selection of the dataset from a monodispersed region for further analysis. The absence of upturned profile at lower momentum transfer (*q*) values in Guinier analysis of SAXS data for *wt* and mutant oligomers implies that all samples are monodispersed and aggregation free (Figure 3A, panel i and ii). Next, the dimensionless Kratky analysis (54) of SAXS data for all samples was performed to assess their flexibility and compactness. This analysis displays a Gaussian curve suggesting that all samples were folded (Figure 3A, panel iii). The SAXS data were then converted to the electron pair-distance distribution plots ((*P*(*r*) function) (Figure 3A, panel iv) using the GNOM software program(55). The characteristic Gaussian-shaped pattern observed in *P(r)* for the *wt* sample indicates that it adopts a compact structure in solution, whereas the *G1738A, G1748A,* and *G1738/1748A* mutant oligomers, despite the same nucleotide length (23 nucleotides), has an increasingly extended structure in solution (Figure 3A, panel iv). The radius of gyration (*R_g_*) for *wt* was calculated to be 18.37 Å, while the *G1738A, G1748A,* and *G1738/1748A* mutants yields increasing values of 19.10, 20.10 and 20.34 Å, respectively, which are in agreement with those obtained from the Guinier analysis (Table 1). The *P*(*r*) function also allows the determination of a maximum particle dimension (*D_max_*) of macromolecules. Based on this analysis, we obtained the *D_max_* of 46.85 Å for the *wt*, while the *G1738A*, *G1748A,* and *G1738/1748A* mutants were 53.14 Å, 60.00 Å and 60.44 Å respectively, indicating that each mutation leads to a degree of alteration of the G-quadruplex structure (Figure 3). To further investigate the structural differences between the samples, we used the *P*(*r*) data in the DAMMIF program that allows low-resolution structure determination. We calculated ten independent low-resolution structures for each sample that provided *X* values ranging from 1.2 to 1.8 indicating the good quality of our models (Table 1). Subsequently, we averaged and filtered ten models using DAMAVER program to obtain a representative low- resolution structure for each sample. We obtained the normalized spatial discrepancy (NSD) values of 0.55, 0.58, 0.58 and 0.59 for *wt, G1738A*, *G1748A,* and *G1738/1748A* mutant samples, respectively, indicating that the ten independent low-resolution structures are highly similar to each other in all cases. The low-resolution structures presented in Figure 3B demonstrate the compact quadruplex structure of the *wt* sample, while each of the mutations displays a relatively extended structure in solution. It is noteworthy that the increase in the size of mutants is also consistent with observed elution volumes from SEC where we observed that the *wt* displayed the highest elution volume (Figure 2A) which progressively increased in the same order as the increase in D_max_ i.e. *wt* < *G1738A* < *G1748A* < *G1738/1748A*.

**Figure 3.**
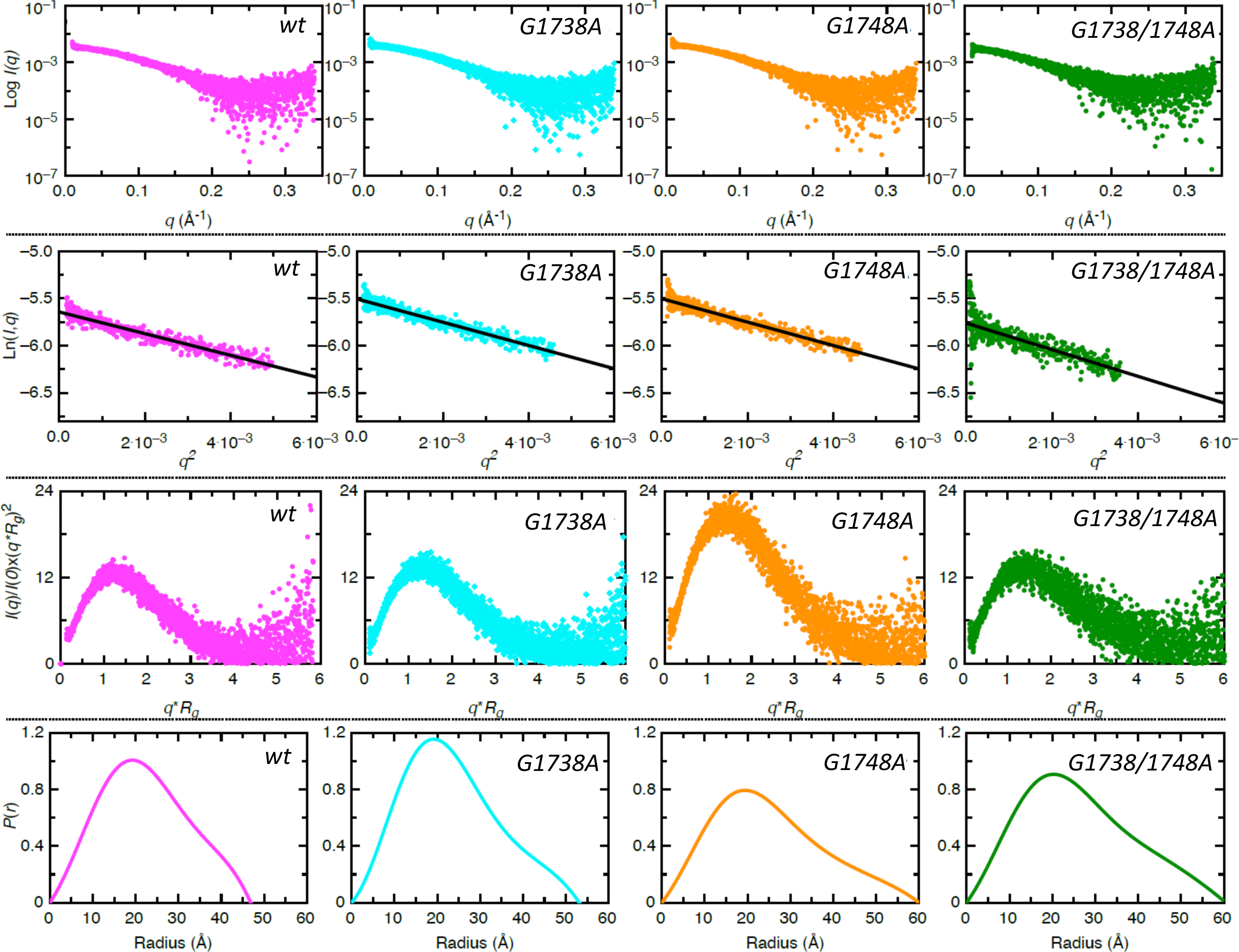
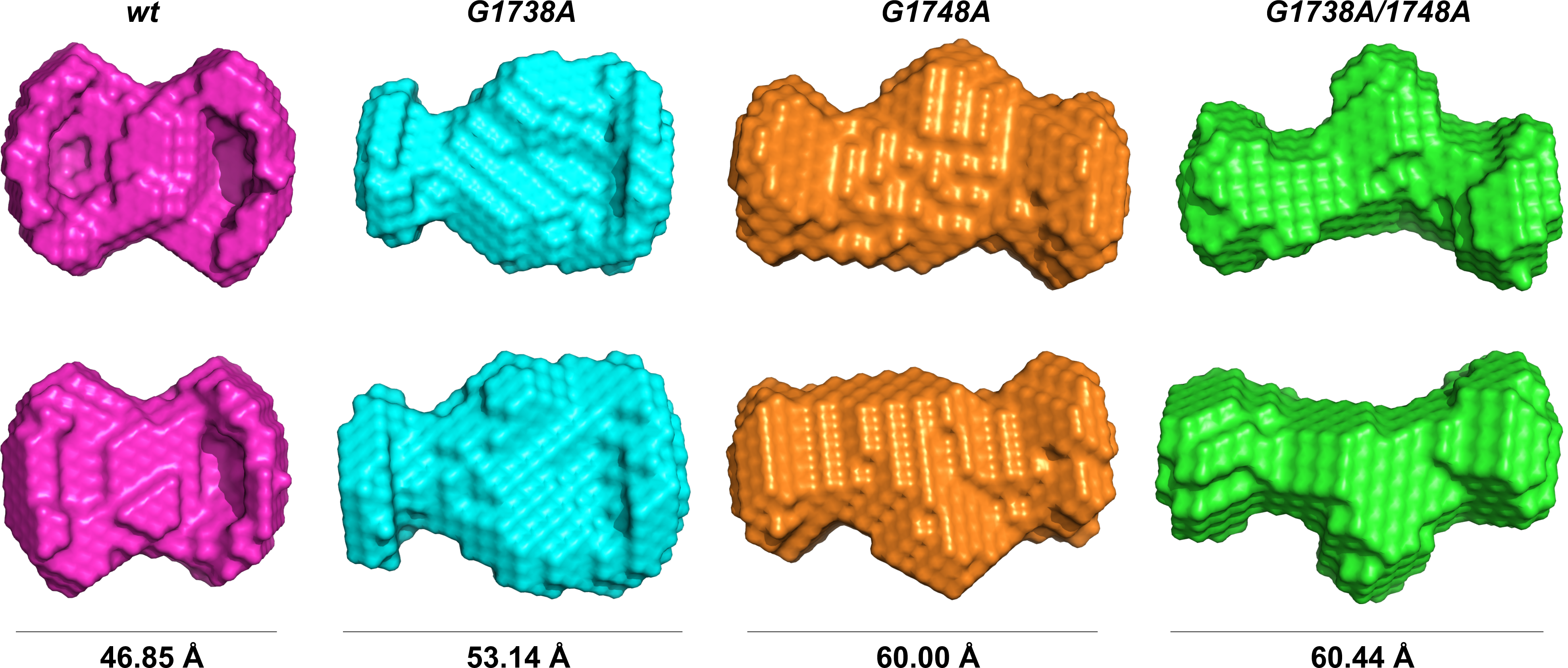
Low-resolution structural studies of HBV pre-core G-quadruplex (wt, pink) and its mutant oligomers (*G1738A*, cyan; *G1748A*, orange and *G1738/1748A*, green). (A, top panel) The top row represents raw SAXS data, where x-axis and y-axis presents momentum transfer (q) and the intensity of scattered light, respectively. The second row indicates Guinier plots *(Ln(I,q)) vs q^2^*) suggesting that all samples are pure. Guinier analysis also provides the *R_g_* for each sample based on the low-q region (See Table1). The third row presenting dimensionless Kratky plots for each sample demonstrates that all samples are folded and suitable for low-resolution model building. Finally, the last row represents the electron pair-distance distribution function plots that provides information about the shape of these molecules, their *R_g_* and maximum particle dimension. (B, bottom panel) Averaged-filtered models derived using DAMMIN and DAMAVER calculations for wt (pink), *G1738A* (cyan), *G1748A* (orange) and *G1738/1748A* (green). The two panels show orthogonal views of G4s, the maximum dimensions (*D_max_*) of the G4 are indicated under each structure, detailed dimensions and parameters are tabulated in Table 1.

To further confirm that the *wt* oligomer forms a G-quadruplex structure, we used the previously described DHX36 protein, (a known G-quadruplex-interacting protein that binds with high affinity to parallel quadruplexes) (53,56–58), and tested for DHX36 ability to recognize the *wt* and/or mutant oligomers via MST. Based on the MST results (Figure 4), we obtained a *K_d_* of 76.81 ± 12.21 nM between DHX36 and the *wt* oligomer further confirming that the *wt* oligomer adopts a G-quadruplex structure in solution that is recognised by DHX36. The *K_d_* values obtained for the mutant oligomers were 14.17 ± 12.76, 132.14 ± 14.84, and 65.50 ± 7.74 nM for *G1738A*, *G1748A* and *G1738/1748A*, respectively.

**Figure 4.**
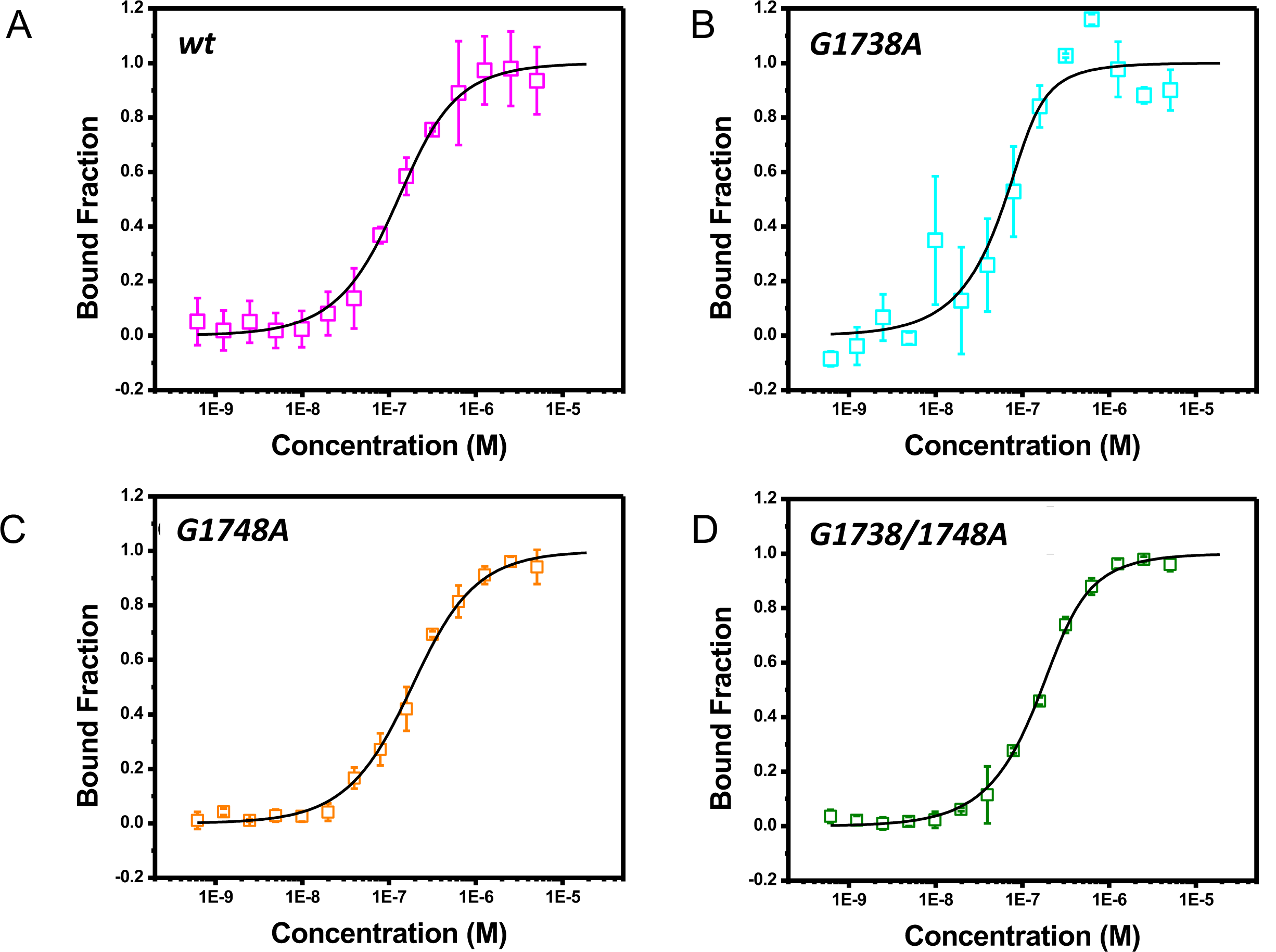
Microscale thermophoresis (MST) of FITC-labelled G-quadruplex oligomers—(A) *wt*, (B) *G1738A*, (C) *G1748A* and (D) *G1738/1748A*—with known G-quadruplex binder, DHX36. The labelled oligomers were held constant while the DHX36 varied in concentration from 0.006 nM to 20 µM in MST buffer (G-quadruplex buffer supplemented with 0.1% Tween20). MST was measured in triplicate and background corrected against spectra of buffer alone. The data above is the average of three independent replicates. Dissociation constants, (K_d_), were computed using K_d_ fitting and are tabulated in Table 2. The binding affinity was observed in the order of *G1738A*<<*wt*<*G1748A*<*G1738/1748A*.

**Table 2:** The dissociation constants calculated for DHX36 53-105 and G-quadruplex variants

| Oligomer | $K_d$ (nM) |
| --- | --- |
| *wt* | $76.81 \pm 12.21$ |
| *G1738A* | $14.17 \pm 12.76$ |
| *G1748A* | $132.14 \pm 14.84$ |
| *G1738/1748A* | $65.50 \pm 7.74$ |

\*The error represents standard deviation of 3 independent experiments.

### The quadruplex-binding protein, DHX36, can pull-down HBV cccDNA derived from HBV infected human liver tissue

The biophysical analysis demonstrated that the *wt* oligomer exists as a G-quadruplex *in vitro*. To address the question of whether the G-quadruplex also exists in HBV genome, we performed a pull-down assay using DHX36-bound-magnetic beads to study if this host protein can bind with HBV cccDNA within HBV-positive human liver tissue. We used nested PCR to amplify HBV DNA in the pull-down products and identified the excess DNA (XS) (*i*.*e*., the initial viral DNA input), but none after the washes (Figure 5B). We then noted a strong HBV DNA signal in the eluted product (EI), demonstrating that DHX36 specifically binds and pull-down HBV cccDNA (Figure 5B). To determine whether DHX36 recognizes other segments of the cccDNA outside the region of interest, two HBV gene segments were amplified (i.e., the C gene with only part of its pre-core promoter region and the X gene, where the 3′ quadruplex-forming region of interest; Figure 5A and S2). The pulldown assay, PCR and gel visualization performed for both of these genes demonstrated an HBV DNA signal (specific PCR amplicon) in the eluted product for the X which contains the G-quadruplex (Figure 5C, EI), but not for the C gene segment (Figure 5D, EI). Mutant X genes show binding with DHX36 in this model as well (Figure 5E), consistent with our findings from MST (Figure 5E). Overall, these results support the presence of the G-quadruplex in the pre-core promoter region of HBV cccDNA.

**Figure 5:**
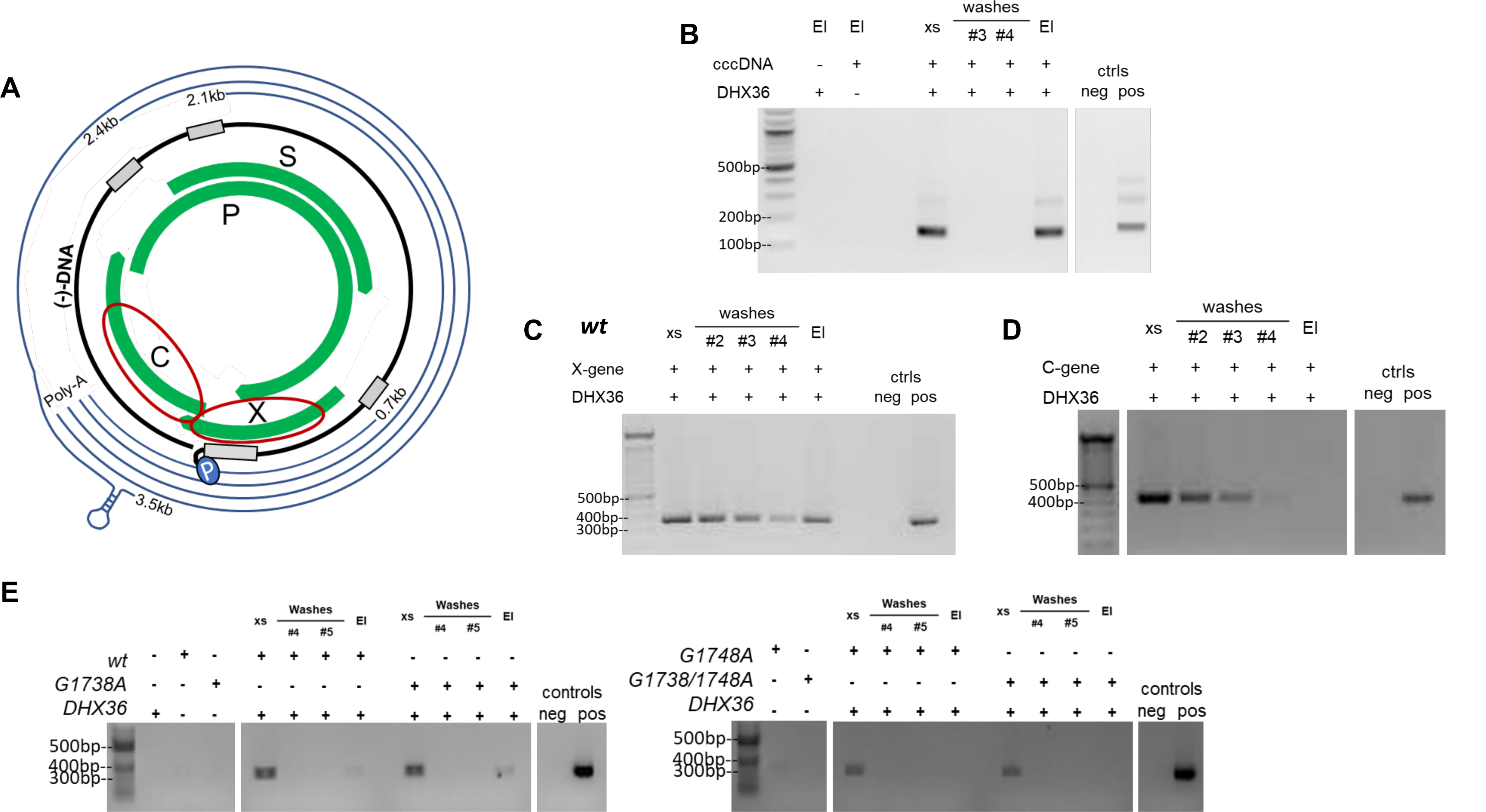
DHX36 pulldown assays. (A) segments of the HBV cccDNA PCR amplified for use in the assay and whole cccDNA extracted from HBV-positive liver tissue; (B) pulldown of whole cccDNA (nested PCR amplified with primers specific to the region of incomplete dsDNA of HBV, nt 1778-1920); (C), pulldown of the X-gene segment; (D), pulldown of the C-gene segment. All PCR were performed on 1:100 dilutions of pull-down washes and elution products. PCR water was used for the negative control; a tandem dimer HBV plasmid was used for positive control. El, Elution; xs, excess DNA. (E) Pulldown study using DHX36 bait protein to bind X gene fragments from the mutant plasmid variants. Sample dilution of 1:800 (left panel) and 1:3200 (right panel), done to account for the slightly differing starting amounts of DNA and the dilution at which the signal is no longer picked up from the nickel bead controls.

### The G-quadruplex region influences HBV replication

Following identification of the G-quadruplex in cccDNA, we investigated the functional impact of this structure on HBV replication. Therefore, we employed a wildtype 1.3mer HBV genome plasmid (kindly gifted from Prof. H. Guo)(59) and designed mutants based on the *G1738A, G1748A,* and *G1738/1748A* sequences used above. The mutant plasmids were transfected into HepG2 cells and a separate GFP control plasmid was simultaneously transfected in all wells to monitor for relative transfection efficiency, inferring relative equivalence (Figure S3). Transfection was consistent across all wells, with an estimated efficiency of about 10% by day 1, and up to 20% for the HBV transfected wells.

The HepG2 transfection studies were performed in triplicate and consistently demonstrated a difference in the viral protein products for *wt* and mutant pre-core promoter regions (Figures 6A-C). The level of HBsAg in the supernatant of HepG2 cells transfected with the *G1738A* and *G1738/1748A* mutant HBV were significantly higher than the *wt* (p<0.05) on days 5 and 7 (Figure 6A, Table S4). Similarly, the levels of HBeAg in supernatant also showed significantly lower production among the wild-type vs. mutant plasmids for nearly all paired sets on days 3 through 7 (Figure 6B, Table S4). In contrast, the high amounts of cellular HBcAg was produced by the *wt* HBV compared to the mutant, with significance noted only between the *wt* and *G1738/1748A* mutant combinations on days 3 through 7 (Figure 6C, S Table S4).

**Figure 6:**
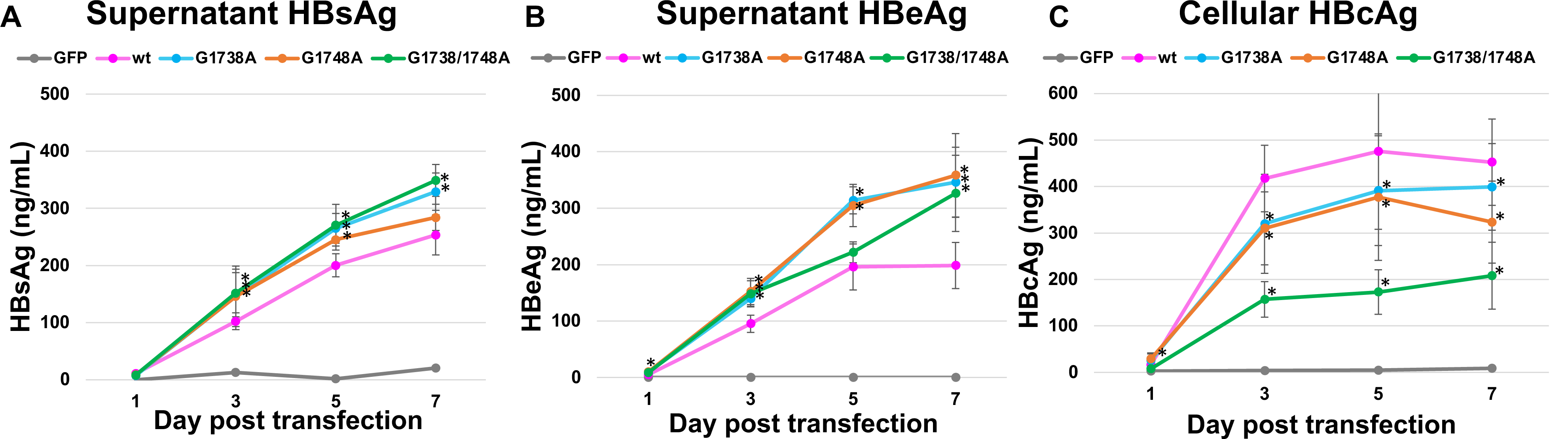
Viral protein markers from 1.3mer HBV transfection studies in HepG2 cells. (A) Average supernatant HBsAg; (B) average supernatant HBeAg; and (C) average cellular HBcAg. Samples were collected at Days 1, 3, 5 and 7 post-transfection and analyzed by various ELISA assays (see Methods). Plots represent the averaged results of the three transfections, with each data point representing at least 8 wells of an ELISA plate. Error bars represent 2x standard error. Significance was calculated using one-way ANOVA with a 2-sided Dunnett t post-hoc analysis. *p-value for significance <0.05. P-values are recorded in the Supplementary Information (Table S4).

Analysis of the nucleic acid levels demonstrated an increase in the amount of cellular DNA detectable above plasmid baseline for all the plasmids at day 7, however, there were no significant differences noted between the wild-type vs. mutant variants after corrections for cell and plasmid (Figure 7, Figure S4-S5). In the supernatant-extracted DNA, a signal was noted on day 7 in one of the three replicate experiments for the *wt* only (Figure 7). The lack of this similar finding in the other replicates likely indicates a lower amount of DNA released into the supernatant relative to the starting plasmid, rather than an experimentally relevant finding. The levels of HBV RNA studies in the supernatant were undetectable when corrected for residual DNA (*i*.*e*., reverse transcriptase negative controls) (Figure 8A; Figure S6). There was a consistent baseline signal in cellular RNA throughout the replicates but no significant difference between the *wt* and various HBV mutants (Figure 8B). Of note, the amount of detectable plasmid decreased over time (by ∼1 log), consistent with the cellular breakdown of the plasmid (Figure S5) thus the detected surplus DNA was derived from the newly replicating virus.

**Figure 7:**
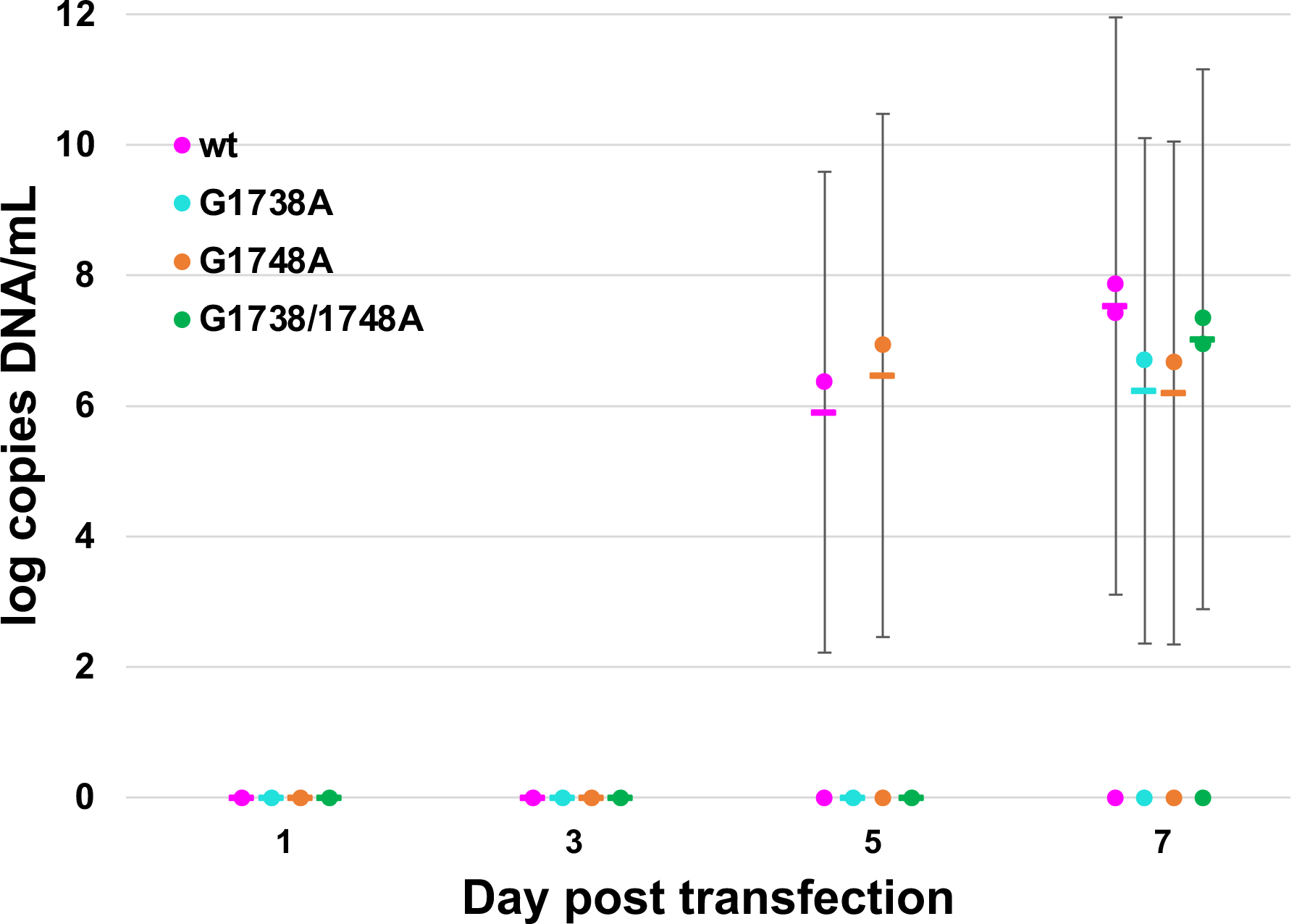
DNA from HepG2 transfection studies with wt and mutant HBV plasmids (*G1738A*, *G1748A*, and *G1738A/1748A*), harvested at days 1, 3, 5, and 7 post-transfection, isolated from the cells. DNA was extracted using phenol-chloroform extraction method from both the supernatant and cells. qPCR was performed using pre-core promoter primers and corrected for plasmid and internal GAPDH control (See Supplementary Information Table S3 and Figures S4- S5). Supernatant DNA was also isolated and quantified, but found in only one replicate (see text).

**Figure 8:**
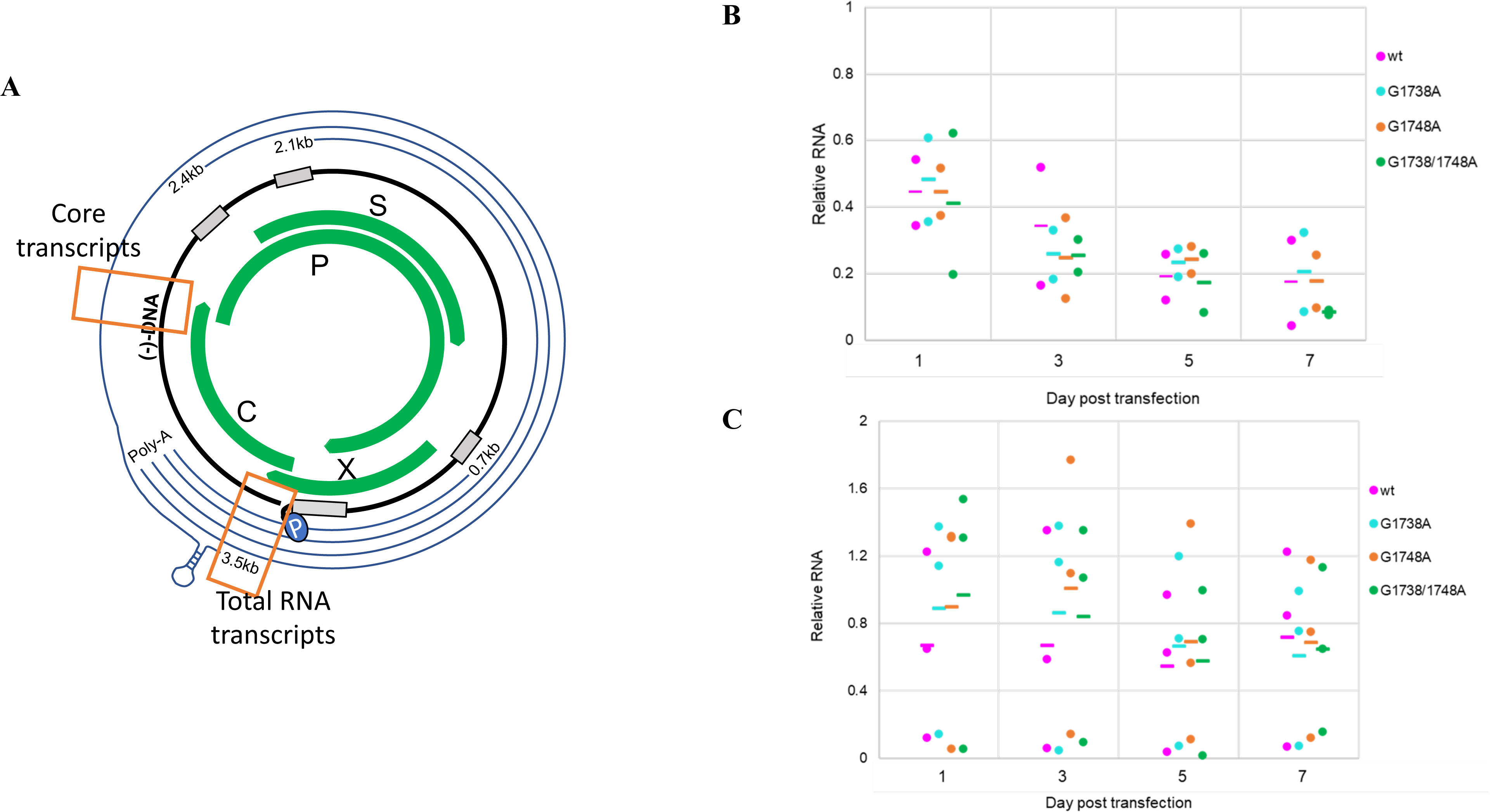
RNA from HepG2 transfection studies with wt and mutant HBV plasmids (*G1738A*, *G1748A*, and *G1738A/1748A*), harvested at days 1, 3, 5, and 7 post-transfection. RNA wa extracted using Trizol extraction followed by DNase digestion. qPCR was performed using primer sets to pick up the core transcripts and total RNA (see Table S3) as noted in the schematic (A). The information is represented as a ratio of core transcripts to total RNA for both the supernatant (B) and cells (C) to account for equivalent detection of residual forms of DNA a detected by qPCR.

## DISCUSSION

There is increasing recognition of G-quadruplexes throughout the human genome (25,26,30,31,47). They are frequently found in key regulatory regions of genes and regulat numerous critical cellular processes including transcription, translation and genome stability (25-28,30,47,60-62). The ubiquity of G-quadruplexes extends to the field of virology where G-quadruplexes have been found in regulatory regions of various viral genomes (36–46).

In this work, we demonstrate the presence of a G-quadruplex in the HBV pre-core promoter region, a region critical to the transcription of the HBV C gene and translation of viral core proteins. The current study describes its evolutionary persistence with the aid of an HBV sequence repository (https://hbvdb.ibcp.fr), which contains an extensive repository of *replication-competent* viruses (63). Across over 10,000 available pre-core/core sequences worldwide, this region is very G-rich and is highly conserved across nearly all major genotypes (Figure 1), except for HBV genotype G (excluded in current analysis). HBV G is rarely found and usually in combination with another genotype (*i*.*e*., A or H), implying a unique strategy to maintain replication (64–66). This strong degree of conservation of this G-rich region provides a rationale for its functional role in virus maintenance, especially in HBV with a highly error-prone polymerase, capable of producing10^10-11^ point mutations daily (67). If a region of the genome were to be of high importance to the virus (and not an inconsequential ‘DNA knot’), it would be evolutionarily maintained over time.

Through rigorous biophysical analysis, we demonstrate the ability of this G-rich HBV pre-core promoter region to form a G-quadruplex, comparing results with those of the HBV single-nucleotide substitutions, *G1738A, G1748A,* and the double mutation *G1738/1748A*. In particular, the *G1748A* mutation has been shown to prevent host Sp1 binding and transcript production (48). We hypothesized that this HBV pre-core mutation affects the quadruplex structure and disrupts host Sp1 quadruplex binding leading to loss of function noted previously (48). Our CD studies support the formation of a parallel quadruplex in the *wt* oligomer as well as the *G1738A* mutant, while the *G1748A* and *G1738/1748A* oligomers form less distinct, likely altered or distorted quadruplexes (Figure 2). This is consistent with reports that CD spectra amplitude differs as a function of G-quartet stacks, thus the altered spectra observed in the case of the mutants support the distortion or alteration of the G-quadruplex (51). More specifically, the *G1738A* mutation does not appear to result in disruption of the “core” of G-quadruplex, likely owing to its more 5′ position, while the *G1748A* mutation does. Further, the SAXS data provide succinct low-resolution structures, indicating the *wt* oligomer indeed forms a quadruplex (*D_max_* = 46.85 Å), while each of the mutants forms an increasingly extended conformation— *G1738A* (*D_max_* = 53.14 Å) > *G1748A* (*D_max_ =* 60.00 Å) > *G1738/1748A* (*D_max_ =* 60.44 Å), highlighting the impact of these mutations on the secondary structure in this region (Figure 3, Table 1). When comparing the effects of the mutations on the resultant secondary structure with these two techniques, the CD and SAXS data agree that the *G1738A* mutant results in a minimal alteration in the G-quadruplex structure. Conversely, the *G1748A* mutation has a greater impact on the structure, which when combined with the *G1738A* mutation, appears to be synergistic in the *G1738/1748A* mutant. Finally, using a known high-affinity parallel quadruplex binder, DHX36, we demonstrate that the *wt* oligomer can interact with DHX36 (Figures 4 and 5), and provide a *K_d_* value for this binding of 76.81 ± 12.21 nM through MST studies (Figure 4). This is a comparable range to other reported *K_d_* values of 0.31 and 0.44 μ binding with a human telomerase RNA G-quadruplex (hTR_1–43_) and basic tetramolecular RNA -A_15_G_5_A_15_ ; ‘rAGA’) (57, 68). This G-quadruplex-binding protein has been shown to interact at picomolar concentrations, whereby full-length DHX36 binds the RNA G-quadruplex ‘rAGA’ with *K_d_* of 39 pM while the DNA equivalent (5′ -3′ pM (69). One of the strongest affinities with DHX36 noted in the literature appears to be with a -untranslated region G-quadruplex structure from the Zic1 gene, a gene responsible for producing a zinc-finger protein critical to early development, with a *K_d_* as low as 3 pM (70). With regards to our mutant oligomers, binding is also demonstrated with *K_d_* values that fall within this broad range. The *G1738A* mutant appeared to have stronger binding than the *wt*, raising the possibility of an alternate binding motif or additional stabilizing interactions with the DHX36 protein. DHX36 has been shown to bind parallel quadruplexes via its N-terminal domain in an end-on manner where its first 21 residues are responsible for the quadruplex binding activity (53,69,71,72). An NMR based study examining the interaction between an 18-amino acid fragment of DHX36 peptide and a parallel quadruplex showed that there are CH/π -stacking interactions occurring between them. This resulted in DHX36 peptide π effectively covering the tetrad from the top, and subsequently positioning the positively charged side-chains of three lysines in closer proximity to the negatively charged tetrads further stabilizing the interaction (72, 73). Data from CD and SAXS experiments demonstrate that all the variants form quadruplexes, hence all of them are capable of binding to DHX36. However, the - and CH3/π -stacking interactions which resulted in the varied strength of *K_d_*. It may also be a feature of a smaller system that could be overcome by employing a longer strand of DNA in which the steric hindrance from a less organized system would disallow binding. A future direction of this work will involve developing a higher-resolution structure through molecular modelling (i.e., molecular dynamic simulations) to create a model consistent with this experimental data.

In addition to *in vitro* biophysical data of controlled oligomers, we demonstrate its presence in HBV positive human liver tissue. We employed pull-down assays using cccDNA extracted from HBV infected liver and demonstrated that the known quadruplex-binder, DHX36 can bind and pull HBV cccDNA from solution (Figure 5B). The inclusion of sub-segments of the HBV genome serves as controls validating the HBV pre-core region as the major binding site. Although the *wt* and *G1748A* mutant X-gene show binding, the *G1738A* and *G1738/1748A* do not bind, and is partially consistent with the structural data analysis. Although it is possible that the contribution of this binding is shared with the other recently identified G-quadruplex in the pre-S region, the prior study did not evaluate DHX36 interactions with the HBV pre-S region (46).

To establish the functional significance of the presence of G-quadruplex in the HBV genome, we compared the wild-type HBV with *G1738A*, *G1748A* and *G1738/1748A* mutant plasmids in *in vitro* cell culture studies. It was postulated that mutations which disrupt the quadruplex region would have reduced binding from host proteins to initiate transcription and thus impact downstream viral replication. To investigate this, we used a wild-type 1.3-mer HBV plasmid(59) and created each of the above mutants (*G1738A*, *G1748A* and *G1738/1748A*) through site-directed mutagenesis. Previous studies with a linearized 1-mer HBV model, in which the HBV genome was excised from the plasmid and allowed to re-circularize by natural DNA repair mechanisms once in the cell, did not prove effective at producing HBeAg or HBcAg at detectable levels (see Supplementary Information —Supplementary Experiments and Figures S7-10)(74). This may have been due to incomplete circularization inside the cell. Since the location of the cut site is through the core gene, the production of HBeAg and HBcAg proteins would be fully reliant on the circularization, whereas the transcripts and open reading frame for HBsAg production would not be affected. Thus, the 1.3-mer HBV model was adopted to overcome these deficiencies, allow for the more balanced production of all viral transcripts and enable an analysis of sequence impacts on viral replication within this cell culture system. We noted differences between mutant and wild-type transfections in some of the markers of viral replication typically after ∼ 5 days (Figures 6-8). Notably, HBsAg in cell supernatant showed that the mutant variants, *G1738A* and *G1738/1748A,* produced significantly more HBsAg than the *wt* on days 5 and 7 (Figure 6A). As the transcript for HBsAg production is independent of the core promoter, the increase in HBsAg for the mutants compared to *wt* promoter region may be explained by a shift in cellular resources. It is possible, that the downregulation of one transcript (*i*.*e*., the core transcript), leads to increased production of other viral transcripts (*i*.*e*., the HBV S transcript). This shift in specific viral protein production could occur if the necessary secondary structure required for transcription factor binding is missing or distorted, as might be the case for pre-core promoter region G-quadruplex in the presence of the *G1748A* mutant. This phenomenon was also observed by Li and Ou (48), where they noted a three-fold increase in S RNA transcript with a G1736 deletion mutation in the pre-core promoter region. Analysis of the HBeAg showed greater levels produced by HBV mutants compared to *wt* virus (Figure 6B). In comparison, a significant increase in levels of HBcAg by the *wt* virus compared to the *G1738/1748A* mutant was observed (Figure 6C). This suggests subtle differences in viral core protein production. Although *nt* changes within each mutant may disrupt the quadruplex, they do not abolish HBcAg production, as would have been expected if the quadruplex was critical to transcription factor binding. This is reflected in our biophysical data where we show the varied binding affinities with the selective quadruplex binder, DHX36. Notably, the double mutation *G1738/1748A* had the weakest binding based on MST analysis and the most unfolded state in the SAXS data and functional studies show it to have the most significant decrease in HBcAg production. The data suggests that even partial quadruplex formation is sufficient for the cell’s usual transcription machinery and any possible subtle differences may not be detectable in the less-efficient cell culture model. Further, the data is consistent with results by Biswas et al., involving the quadruplex study of HBV’s S promoter region in which only a partial decrease in downstream products rather than complete loss of viral protein due to quadruplex-disrupting mutations(46).

We determined the effect of the *G1738A*, *G1748A* and *G1738/1748A* mutations on the production of viral nucleic acid products. The total supernatant DNA was detected on day 7 post-transfection in only one of the three replicates (log 5.7 copies DNA/mL in transfection #2). The cellular DNA was detectable for all plasmid forms on day 7, but without significant differences among the *wt* and mutant comparators (Figure 7). Total HBV RNA in the supernatant and in the cells was at a very low-level when corrected for residual DNA (i.e., starting plasmid or other HBV DNA products). To overcome potential confounders from larger amounts of DNA, we compared the RNA transcripts as a ratio of those produced from the core promoter (*i*.*e*., pre-core and pgRNA transcripts) to the total RNA (*i*.*e*., as measured by a region just prior to the poly-A tail signal) (Figure 8A). We assume the relative PCR efficiency for each of the two primer sets is comparable and equivalent for each comparator thus these values represent a relative impact of the mutation on core and pgRNA transcript production. For *wt* HBV, the ratio of supernatant core and pgRNA transcripts appear to decrease over time, whereas in the cells, this amount appears to be relatively stable (Figure 8B,C). For the mutant HBV plasmids, the ratios of core to total RNA transcripts follow a similar pattern, without a clear trend for favouring alternate transcripts.

Overall, using the 1.3-mer HBV plasmid model, the differences in the *wt* and various mutant pre-core promoter HBV transfections suggest the only minimal influence of the mutations on overall downstream function. There are several possible explanations for this finding; (1) transcription factor binding to promoter regions of the 1.3-mer HBV plasmid is independent of or only partially dependent on the quadruplex under study. It is possible that in the setting of high levels of plasmid, even a less-optimal binding alternative may still favour production despite potential structural controls. Therefore, this model may not be representative of natural processes despite successes with this approach for other pathway studies (59,75,76). (2) It is also possible that the quadruplex structure is not needed for binding and that the primary nucleotide sequence is the sole influencer of the subtle finding in this model. Fully delineating the two possibilities in an experimental model would require conditions that affect the quadruplex formation *in vivo* via the stabilizing metal ion, and that would still be compatible with cell growth or function. While known small molecule G-quadruplex stabilizers would be another possibility to investigate this phenomenon, the off-target effects of these agents on the cells’ normal functioning may limit evaluation.

In summary, our work provides robust biophysical evidence and novel functional data for the existence of a G-quadruplex structure in the pre-core promoter region of the HBV genome. The data demonstrates that we can use a known quadruplex-binder to bind a physiologically-relevant, transcriptional template of HBV—the resilient cccDNA. Furthermore, using a well-validated HBV culture model, we provide evidence for a functional effect on HBV replication.

This unique structural motif represents a novel HBV therapeutic target the HBV cccDNA minichromosome and potentially informs the development of a sterilizing or virological cure for HBV infection.

## EXPERIMENTAL PROCEDURE

### Analysis of the HBV core promoter region

The HBV genome database (HBVdB: https://hbvdb.ibcp.fr, accessed April, 2019) was used to access 9939 HBV core promoter sequences from 7 of the8 main HBV genotypes (977 genotype A; 2478 genotype B; 3313 genotype C; 1545 genotype D; 389 genotype E; 387 genotype F; and 52 genotype H). Sequences were downloaded as a Clustal W alignment (77), truncated as the 23-mer region (nt1732-1754) and examined for the presence and frequency of single nucleotide polymorphisms with the aid of the Geneious Prime program (2019.1.3; www.geneious.com). HBV genotype G sequences lack the highly G-rich region and were excluded from analysis (21).

### Quadruplex preparation

All DNA oligomers were ordered from AlphaDNA (Montreal, QC, Canada), with high-performance liquid chromatography (HPLC) purification by the manufacturer. The pre-core

-CTGGGA**G**GAGCTGGGG**G**AGGAGA-3′ and

-CTGGGA**<u>A</u>**GAGCTGGGG**G**AGGAGA-3′,

-CTGGGA**G**GAGCTGGGG**<u>A</u>**AGGAGA-3′ and (*G1738/1748A*) 5′

. Samples were dissolved in G-quadruplex buffer (20 mM HEPES, pH 7.5, 100 mM KCl, 1 mM EDTA) at a concentration of 5 μ spectrophotometrically using extinction coefficients for 260 nm (calculated using IDT OligoAnalyzer tool) of 237,400 M^-1^ cm^-1^(*wt*), 239,300 M^-1^ cm^-1^(*G1738A* and *G1748A*) and 241200 M^-1^ cm^-1^ (*G1738/1748A*). Samples were heated to 95°C for ten minutes, then slowly cooled to room temperature to allow assembly of the G-quadruplex. Samples were concentrated L using a 3 kDa Vivaspin™ 20 filter column (GE Healthcare, Mississauga, ON, μ Canada). The conformational mixtures were purified via size exclusion chromatography (SEC) using a HiLoad Superdex 75 10/300 in G-quadruplex buffer, loaded with 500 μ eluted at 0.65 mL/min using G-buffer (20 mM HEPES, pH 7.5, 100 mM KCl, 1 mM EDTA). Only peak fractions (e.g., for G1738A, only fractions from ∼12.5 to 15 mL, see Figure 2A) were pooled and concentrated using a 3 kDa spin column, for use in subsequent experiments.

### Circular Dichroism (CD) spectropolarimetry

All spectra for wt and mutant G-quadruplexes were recorded on a calibrated Jasco J-815 spectropolarimeter (Jasco Inc., Easton, MD) from 220–320 nm in a 1.0 mm cell and a 32 s integration time. SEC purified oligomers were concentrated to 20 μ Measurements were performed in triplicate and baseline-corrected by subtraction of the degassed buffer alone.

### Molecular cloning and protein preparation of DHX36

DHX36 was prepared as previously described (78). Briefly, amino acid residues 53-105 of human DHX36 were PCR amplified from a gBlock® (Integrated DNA Technologies, Kanata, ON, Canada) to add NdeI and SalI restriction sites. The PCR product was inserted into the pET28a(+) vector using NdeI and SalI restriction enzymes and T4 DNA ligase (Thermo Fisher Scientific, Waltham, MA) and subsequently transformed into chemically competent DH5α *E. coli* cells (New England Biolabs (NEB), Ipswich, MA). Positive clones were identified, and the gene sequence was confirmed by Sanger sequencing (Genewiz, South Plainfield, NJ). The final pET28a-DHX36_53-105_ expression vector contained a vector-encoded N-terminal hexahistidine tag and a thrombin cleavage site for tag removal. The pET28a-DHX36_53-105_ expression vector was transformed into competent *E. coli* cells Lemo21(DE3) (NEB), and expression induced via the -D-1-thiogalactopyranoside. Cells were harvested three hours post-β induction by centrifugation at 5000 g for 15 minutes, flash-frozen, and stored at -80°C. Purification of the protein was performed via a nickel affinity column, followed by SEC as described previously (52, 53).

### Microscale Thermophoresis (MST)

MST was used to determine the nature and strength of the interactions between the G-quadruplex with DHX36, where the G-quadruplex oligomers were labelled with FITC and held constant, similar to described (79, 80). First, DHX36 was diluted from 20 µM to 0.006 nM in MST buffer (G-quadruplex buffer supplemented with 0.1% Tween20). Next, the folded and SEC-purified oligomers were diluted to 50 nM in MST buffer and added to each of the tubes of the DHX36 dilution series. Samples were run using Monolith NT.115 standard capillaries (Nanotemper Technologies, San Francisco, CA). The data were collected at the high excitation setting with the laser power set at 20%. Microscale thermophoresis was measured in triplicate and background corrected against spectra of buffer alone. The data from three independent replicates were analyzed using MO Affinity Analysis software v2.1.3 and fit to the standard *K_d_* fit model.

### Small angle X-ray scattering (SAXS)

HPLC-SAXS data for the wildtype and mutant variant oligomers were collected using the B21 BioSAXS beamline at the Diamond Light Source (Didcot, UK) synchrotron facility, under a previously described protocol (52). Using an Agilent 1200 (Agilent Technologies, Stockport, UK) in-line HPLC with a specialized flow cell, oligomer data was collected at 100 μ concentrations in G-quadruplex buffer. Each oligomer sample was injected into a Shodex KW403-4F (Showa Denko America, Inc.) column pre-equilibrated in G-quadruplex buffer. X-rays were exposed to each frame for 3s and on average, 10 frames from each individual sample peak were integrated, followed by buffer subtraction, and then merged using ScÅtter (81) as previously described (80).

The data was imported into software Primus from the ATSAS suite of software programs version 2.8 (82), Guinier analysis was performed on the dataset to ascertain the quality of the data and estimate the Radius of Gyration (*R_g_*) (Figure S1). For calculation of *ab initio* low resolution structures the maximum, linear dimension (*D_max_*) was estimated using the pairwise distance distribution function *P(r)* plot, along with the *R_g_*, estimated using GNOM (55). Using the *P(r)* plot information, ten models were calculated using DAMMIN (83), by selecting a different random seed for each model but keeping identical parameters within each set ensuring no enforced symmetry (*P1*). The program DAMAVER (84) was used to average and filter the models to produce a representative model, as described previously (85).

### Isolation and preparation of HBV cccDNA

HBV DNA positive liver tissue was obtained under an approved ethics protocol (University of Calgary Conjoint Health Research Ethics Board, ID# 16636). Protein-free, nuclear DNA was extracted from liver by Hirt extraction method, as previously described (86). Samples were treated with T5 exonuclease (NEB, Ipswich, MA, Cat# M0363) to digest relaxed circular DNA (rcDNA) or remaining genomic fragments. Products were further cleaned by column purification (E.Z.N.A. Cycle-Pure Kit, Omega Bio-tek, Norcross, GA) to remove the exonuclease. HBV cccDNA was amplified by nested PCR, using HBV specific primers spanning the nick region of the rcDNA genome, as described (86, 87).

### Amplification of HBV DNA fragments

A full-length HBV (genotype C) plasmid was used for PCR amplification to produce HBV DNA fragments of the C gene and promoter region that included the quadruplex forming region (X fragment, “X”) or not (C fragment, “C”), using HBV specific gene primers coding for *nt* 1606-1974 (368bp, “X”) and *nt* 1825-2286 (461bp, “C”), as described (87). The PCR amplicons were visualized on a 1.2% agarose gel and purified using the Qiagen gel DNA extraction kit (Qiagen Inc, Montreal) (See Figure S2, Table S3). The gel-purified products were heated to 95^◦^C for 5 mins, then cooled slowly to room temperature to enable folding.

### HBV genome pull-down assays

Magnetic beads (HisPur Ni-NTA, ThermoFisher, Cat# 88831) and magnetic tube rack (Qiagen Inc, Montreal) were used for HBV genome pull-down assays using DHX36 (with an associated His-tag) as the “bait” protein. The protocol provided with the magnetic beads was loosely followed, with modifications for our system. Firstly, 40 μ added to each tube and equilibrated with washes of equilibration/wash buffer (30 mM imidazole in phosphate-buffered saline (PBS), 0.05% Tween, pH 7.5), per protocol. Secondly, a 1:1 solution of DHX36 in equilibration/wash buffer (end protein concentration of 10 μL/tube) on an end-on-end rotor for 30 mins at 4°C. The excess μ protein was removed, and the beads were washed with buffer until fluorometry readings showed undetectable protein for two consecutive readings (four washes). A dilution of cccDNA (whole or fragments) was made with the buffer to provide 400 μ L, respectively, for the whole cccDNA, C and X regions). The viral DNA was μ incubated with beads (+/- bound DHX36) for 2 hours on a rocker at 4°C. Following removal of excess DNA, the beads were washed with buffer until fluorometry readings showed undetectable DNA for two consecutive readings (four washes). The protein-DNA complexes were eluted with L elution buffer (250 mM imidazole in PBS, pH 7.5).

### Detection of cccDNA and cccDNA fragments

To confirm the presence of HBV cccDNA in the Hirt-extracted, T5 exonuclease-digested liver samples, nested PCR was performed to amplify the “nicked” region of the rcDNA, or -ACTCCTGGACTCTCAGCAATG-3′ (cccDNA-D-For); 5′-GTATGGTGAGGTGAGCAATG-3′ (cccDNA-D-Rev); 5′- AGGCTGTAGGCACAAATTGGT-3′ (cccDNA-N-For); and 5′ GCTTATACGGGTCAATGTCCA-3′ (cccDNA-N-Rev). The products obtained from the X and C region pull-downs were PCR-amplified using the same primers used for their production (see above) and amplicons visualized with agarose gels and SafeView® (Applied Biological Materials Inc., Vancouver) staining under UV light.

### HBV plasmids for transfection of HepG2 cells

HBV mutants (*G1738A*, *G1748A* and *G1738/1748A*) were created using a 1.3mer HBV plasmid (gifted by Dr. Haitao Guo)(59) and site-directed mutagenesis kit (QuikChangeII, Agilent Technologies, Santa Clara, CA) with corresponding primers (University of Calgary DNA Synthesis Lab, Calgary, AB) (see Table S2). The mutant HBV sequences were confirmed by Sanger sequencing. The original plasmid (*wt*) and the core-mutants (*G1738A*, *G1748A* and *G1738/1748A*) were amplified via an *E. coli* (Top10) system (Thermo Fisher, Cat# C404010) and isolated using the GenElute Plasmid Miniprep kit (Sigma, Cat# PLN350). In addition, an in-house green fluorescent protein (GFP)-pcDNA3.1(+) plasmid was used as control.

### HepG2 transfections and isolation of total nucleic acid and protein

HepG2 cells (New England Biolabs) were grown to 80-90% confluence in Dulbecco’s Modified Enrichment Media (DMEM) supplemented with 10% fetal bovine serum and 100 U/mL penicillin/streptomycin. Approximately 24 hours prior to transfection, the cells were split and seeded into 6-well plates at concentrations of 1 x 10^6^ cells/well and allowed to re-attach overnight. On day 0, cell media was exchanged to remove any unattached cells. The corresponding wells were transfected with GFP alone, GFP + HBV_wt_ or GFP+HBV_mutant (G1738A, G1748A or G1738/1748A)_ (500 ng GFP plasmid/well and 270 ng HBV plasmid/well), using Lipofectamine3000 (ThermoFisher) per the manufacturer’s protocol. On days 1, 3, 5 and 7, cells were imaged for GFP to determine % transfection relative to GFP signal, and subsequently, both the supernatant and cells were harvested for detection of HBV RNA, total HBV DNA, and/or viral proteins. Total RNA was extracted via TRIzol® (Invitrogen, Carlsbad, CA), as described (87); total DNA was isolated from cells via phenol-chloroform (supernatant) or TRIzol®. Viral proteins (HBsAg, HBeAg) were detected in the cell supernatant without further manipulation and using culture media for controls (see ELISA details below). HBcAg in cells was detected from the lysate of PBS-washed cells treated with 1% Triton X-100 in TBS buffer. All nucleic acid extractions were performed with simultaneous “mock” extractions under strict lab protocols to prevent cross-contamination. Transfection experiments were performed in triplicate with duplicate samples for each condition per day, per experiment.

### Quantification of HBV RNA and DNA

HBV RNA from the transfected cell supernatant was detected by TaqMan® qPCR following the protocols previously described (88). Total supernatant DNA, cellular RNA and DNA were detected using SYBR® Green fluorescence (BioRad, Cat# 1725124). An HBV-containing plasmid was used as a positive control and to produce the dilution series for the qPCR standards; water and “mock extracted” template and mock transfection were used as negative controls. A segment of the pre-core promoter region was amplified using the primers HBV-PCP- For and HBV-PCP-Rev and subsequently corrected for starting plasmid using primers Plasmid- Guo-For and Plasmid-Guo-Rev (See Table S2). Samples were prepared in triplicate with concomitant standards and controls per qPCR run and all transfections were performed in triplicate. The qPCR amplicons from the *wt*, various mutants and GFP transfected controls were confirmed by agarose gel visualization to confirm single products. In the GFP control, a qPCR signal was noted with a different melting curve than the HBV product and this was confirmed to be a single 800-bp product on a gel that was not seen in the HBV transfected samples, where only the expected HBV fragment size of 340 bp was seen. Primer alignment of the GFP pcDNA3.1(+) plasmid confirmed this amplicon size and poor sequence overlap. HBV nucleic acid detection was corrected for each experimental condition (per the GAPDH internal control).

### Detection of HBsAg, HBeAg and HBcAg

HBsAg was detected by sandwich ELISAs via an in-house developed assay. Briefly, using a 96-well plate (Corning Incorporated, Corning, NY, Cat# 3361), ELISA plates were prepared using HBsAg capture Ab (Fitzgerald Industries International, North Acton, MA, Cat#10-H05H) at 1 ug/mL. An HBsAg dilution series (500 ng/mL - 4 ng/mL) was prepared for use as standards (Fitzgerald, Cat #C-CP2019R). To each corresponding well, 100 μ supernatant or standards was added in triplicate and incubated overnight. The HBsAg was detected using an HRP-conjugated anti-HBsAg antibody (Fitzgerald, Cat# 60C-CR2100RX, 1:2000 dilution), activated by TMB (LifeTech Novex, Cat#00-2023), quenched by 1 M HCl and read at 450 nm (BioRad iMark, USA). The total protein per sample was determined via the BCA protein assay standards kit (Biorad, Cat# 500-0006) and used to correct for varying protein content within the cell supernatant. HBeAg in the supernatant was analyzed via an electrochemiluminescent immunoanalyzer (Roche Elecsys run on the Cobas e411 immunoanalyzer, Roche Diagnostics, Laval, QC) using the sandwich principle involving a reaction with two HBeAg-specific monoclonal antibodies (one biotinylated and the other labeled with a ruthenium complex). The complex was bound to the solid phase consisting of streptavidin-coated magnetic microparticles (assay detection limit of is ≤ 0.30 Paul Ehrlich Institute U/mL, corresponding to 1.6 cut-off index (COI) in the assay’s detection units). Additionally, cellular HBcAg was analyzed using a commercial kit (Cell Biolabs Inc, San Diego, CA, Cat# VPK150), following the manufacturer’s instructions. All data shown is an average of three independent runs performed as duplicates.

### Statistical analysis

Data were analyzed using SPSS statistical software, ver 26 (SPSS Inc., Chicago, IL, USA) and Microsoft Excel. One-way ANOVA analyses with a 2-sided Dunnett t post-hoc analysis were employed to determine the significance of the transfection parameters. Statistical significance was set at *p* values of less than 0.05.

### DATA AVAILABILITY

All data has been presented in this article.

## ACKNOWLEDGEMENTS

We thank Dr Hans-Joachim Wieden for allowing us to use the circular dichroism spectroscopy instrument.

## ACKNOWLEDGEMENT

TRP & CC: conceptualization of the study, designing of experiments, analysis of data, preparation of manuscript, VMS: designed experiments, performed molecular and cell biology experiments analysis of data, preparation of manuscript, MDB: designed experiments, performed biophysics experiments analysis of data, preparation of manuscript, TM: performed MST experiments, preparation of manuscript KCKL: assisted in cell biology experiments, preparation of manuscript SKS: designing of DHX36_53-105_, DLG: expression and purification of DHX36_53-105_ for cell biology experiments, CO: performed the quantitative ELISA experiments, GvM: data analysis. All authors contributed to manuscript preparation.

## SUPPLEMENTARY INFORMATION

Supplementary information is available at JBC online.

## ABBREVIATION

HBV: Hepatitis B virus
*R_g_*: Radius of Gyration
*D_max_*: maximum dimension
MST: Microscale Thermophoresis
SAXS: Small-angle X-ray Scattering

