## Supplemental data for "Identification and characterization of a G-quadruplex structure in the pre-core promoter region of hepatitis B virus"

\# Equal author contribution

**SUPPLEMENTARY FIGURES**

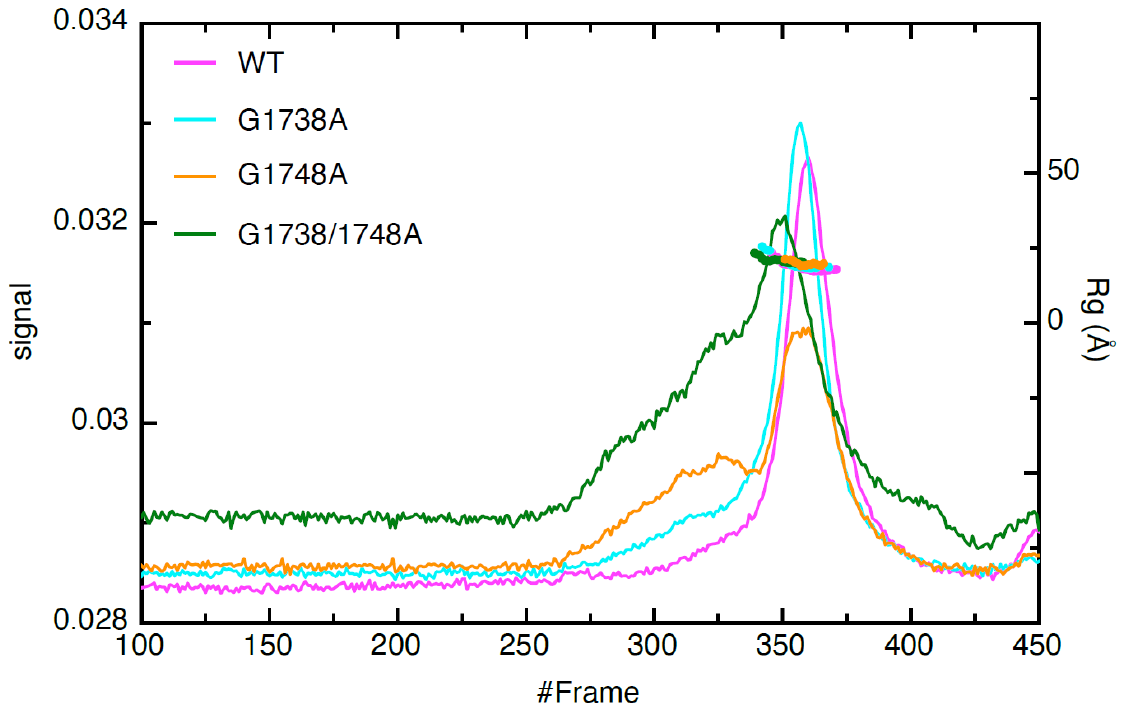

**Figure S1** - HPLC-SAXS profile for HBV pre-core G-quadruplex. The X-axis represents frame numbers, whereas the y-axis on the right indicates the distribution of Rg for each sample. The y-axis on left represents the scattering signal.

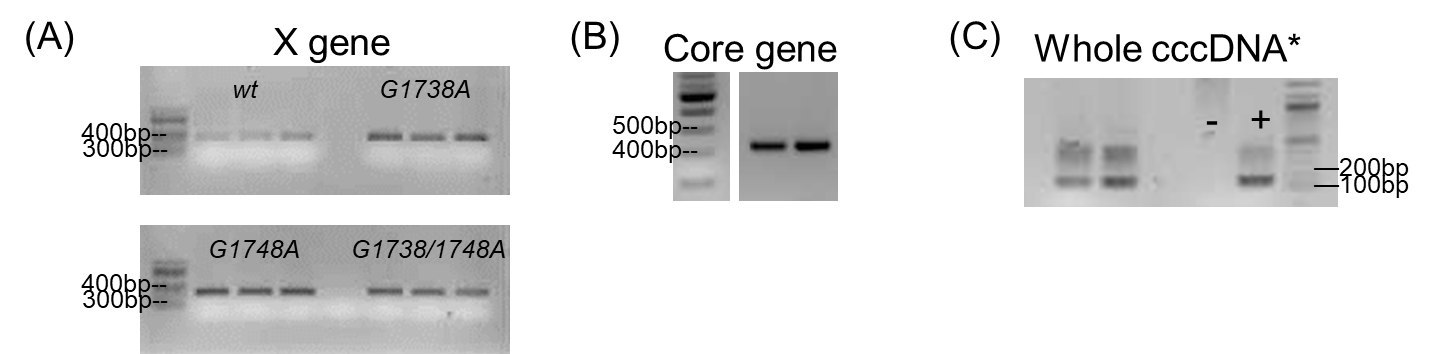

**Figure S2:** Gel electrophoresis of HBV DNA fragments. (**A**) HBV “X” gene fragments from *wt*, *G1738A*, *G1748A* and *G1738/1748A*; (**B**) HBV “C” gene fragment; and (**C**) Whole cccDNA extracted from liver tissue, after PCR amplification with HBV specific primers spanning the nick region of the HBV rcDNA genome.

*Amplified nicked region: nt 1778-1920; positive control is an in-house HBV-dimer plasmid.

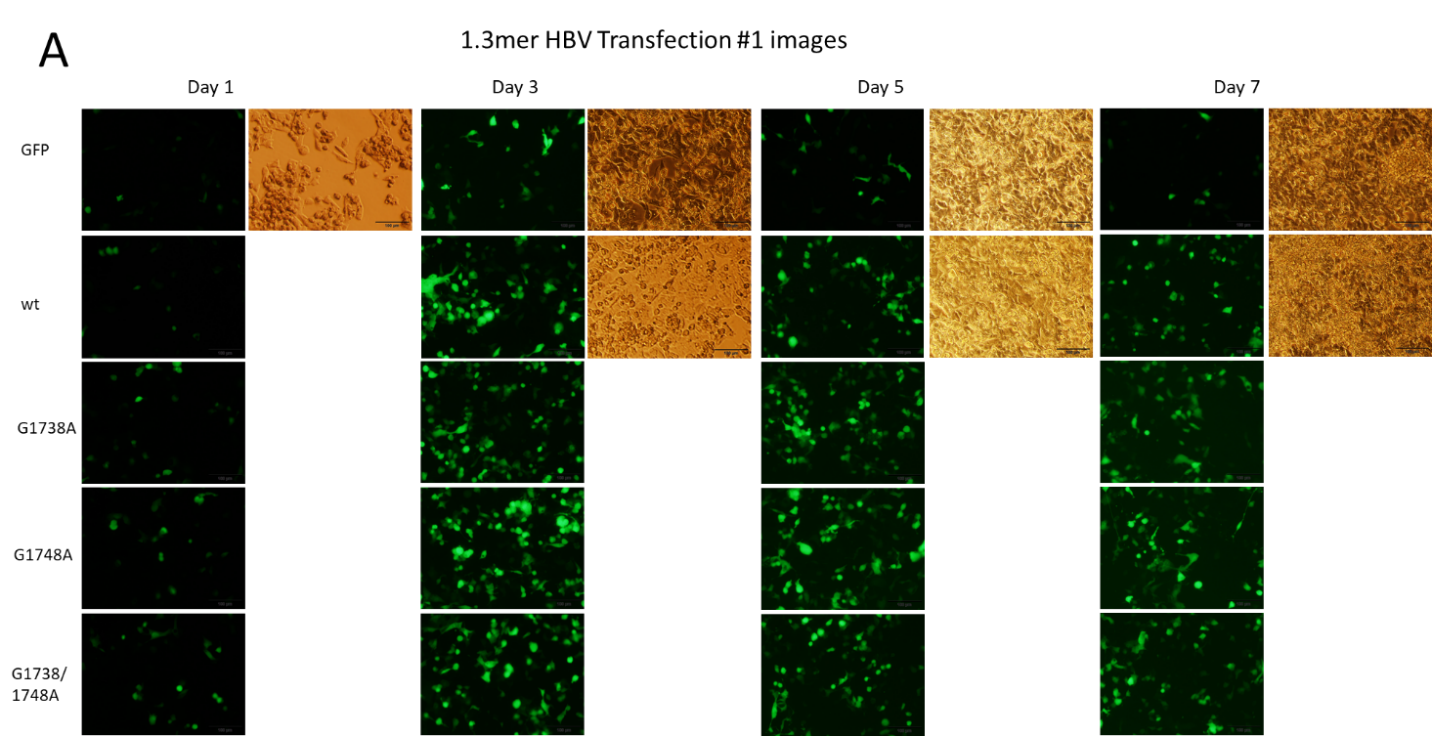

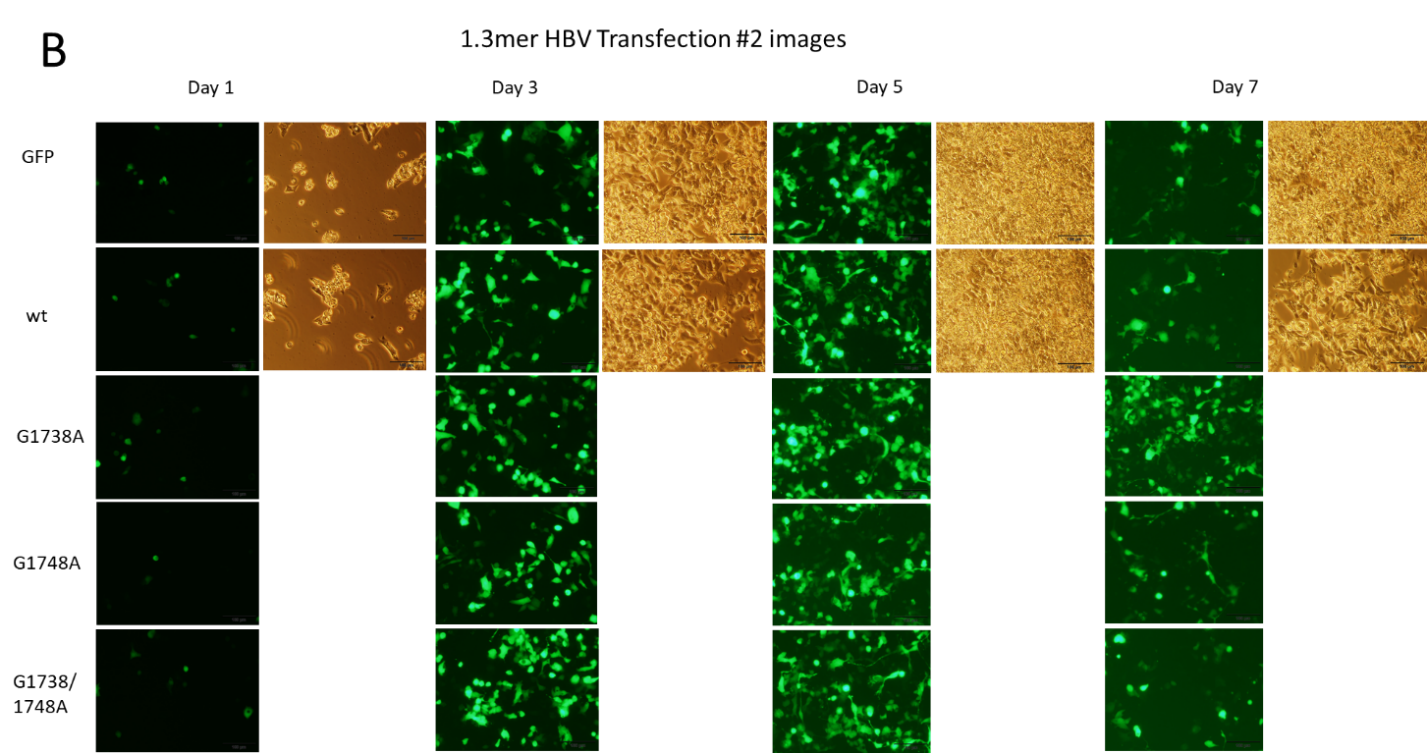

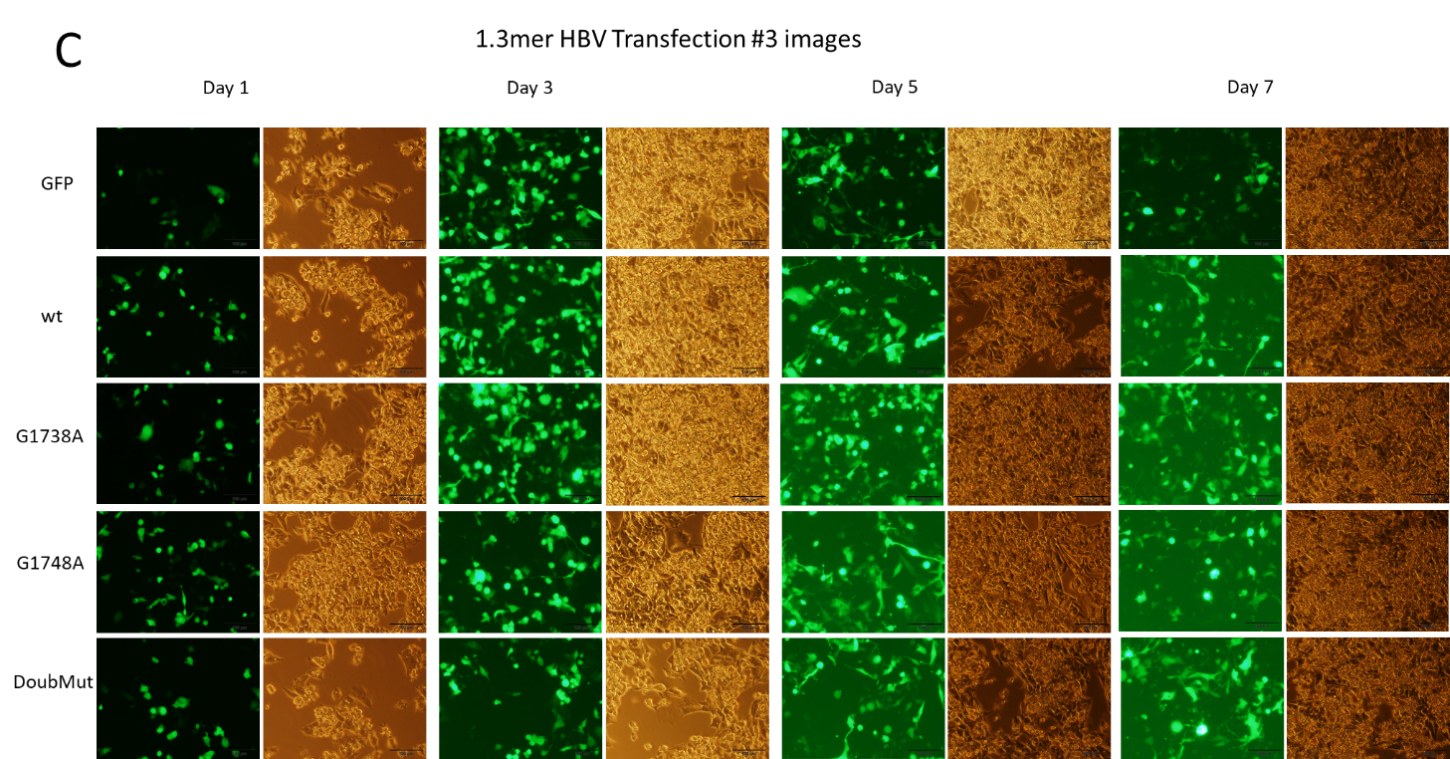

**Figure S3**: Images of 1.3-mer HBV transfection of HepG2 cells taken on days 1, 3, 5, and 7, for each Transfection—(A) #1, (B) #2, and (C) #3. For the *wt* and GFP controls, phase contrast images were taken for all for comparison. For those in which phase contrast images were not taken, such as the mutants of Transfections #1 and #2, visual inspection at the time revealed a similar cellular density to that of the *wt* wells.

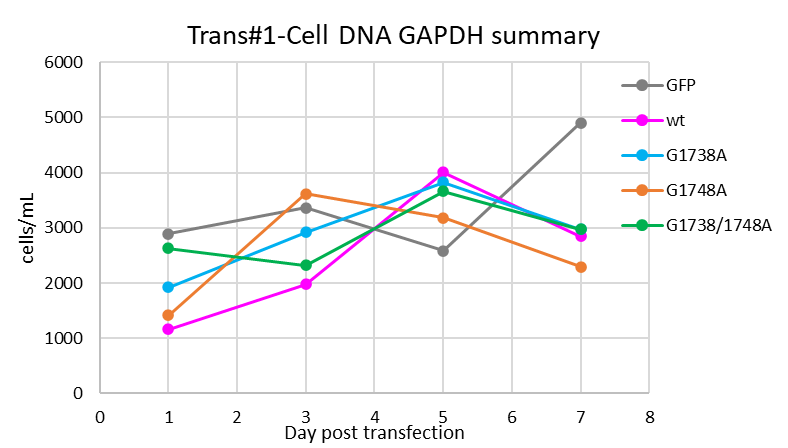

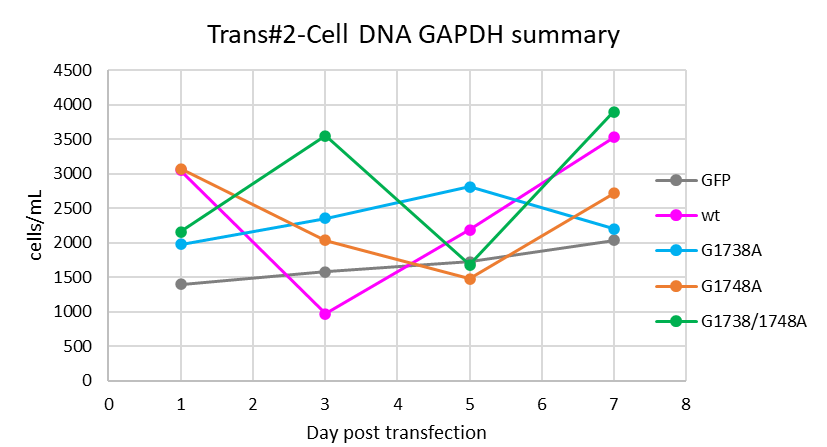

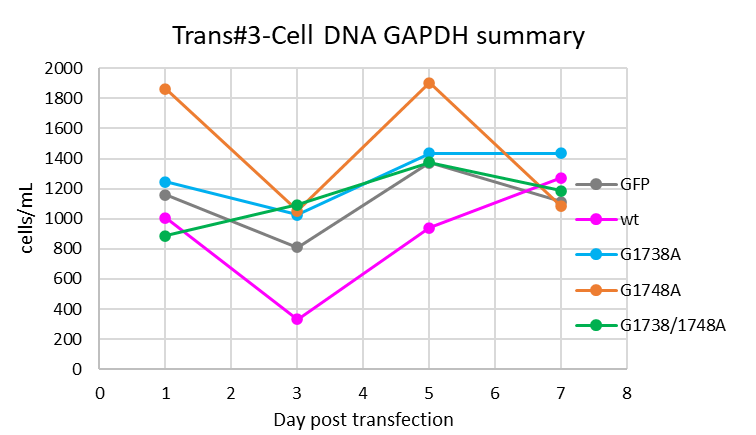

**Figure S4**: Cellular GAPDH from each of the transfection studies as measured by qPCR of the products from DNA extraction.

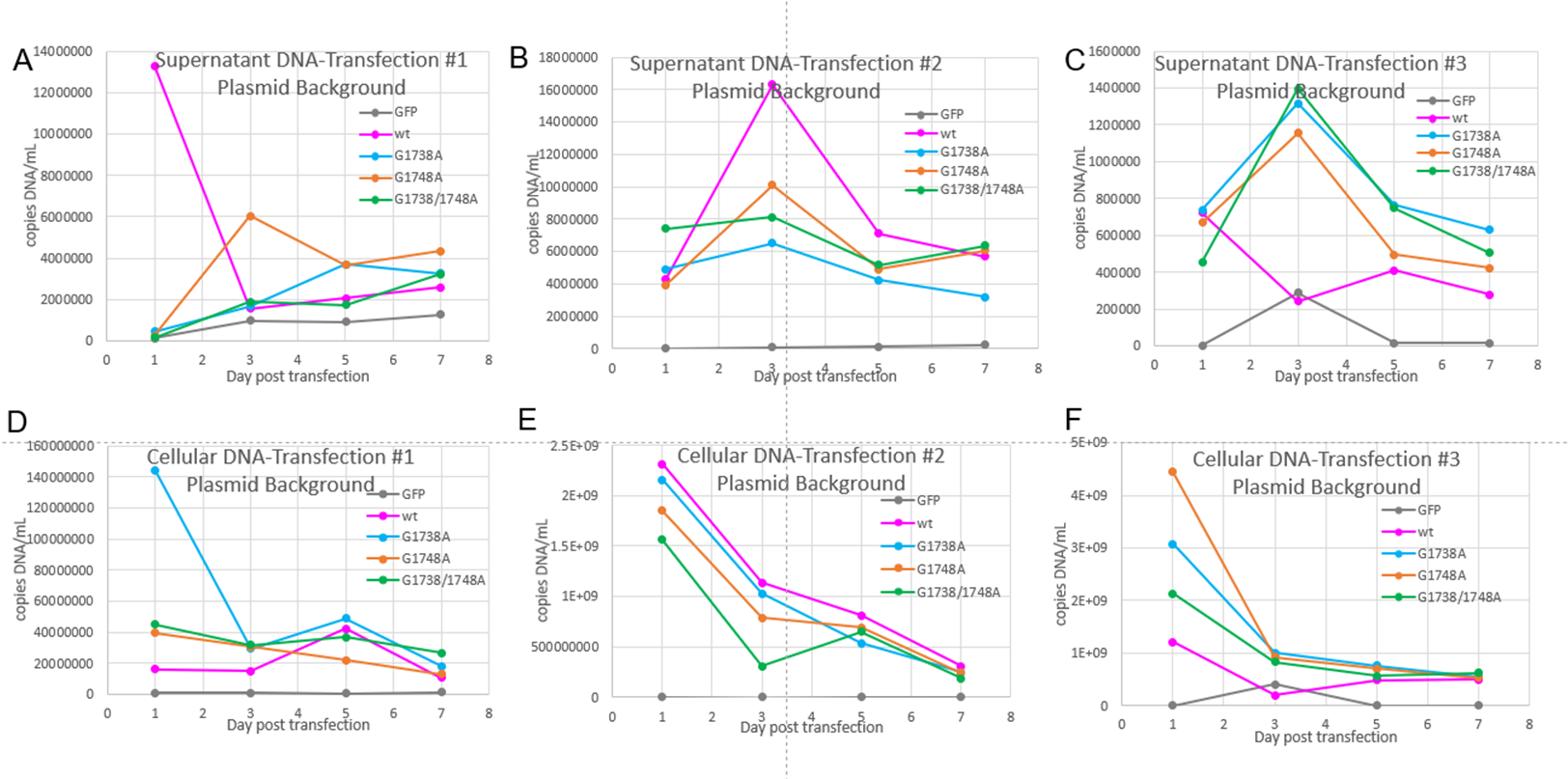

**Figure S5**: Plasmid detection in the supernatant (A-C) and cells (D-F) of the transfection experiments. Of note, the cellular DNA of transfection #1 (D) was extracted using the Hirt extraction method, whereas transfections #2 and #3 (E and F) were extracted using standard DNA extraction methods. Also of note is the plasmid decay within the cells, particularly seen in Transfections #2 and #3 extractions (E and F).

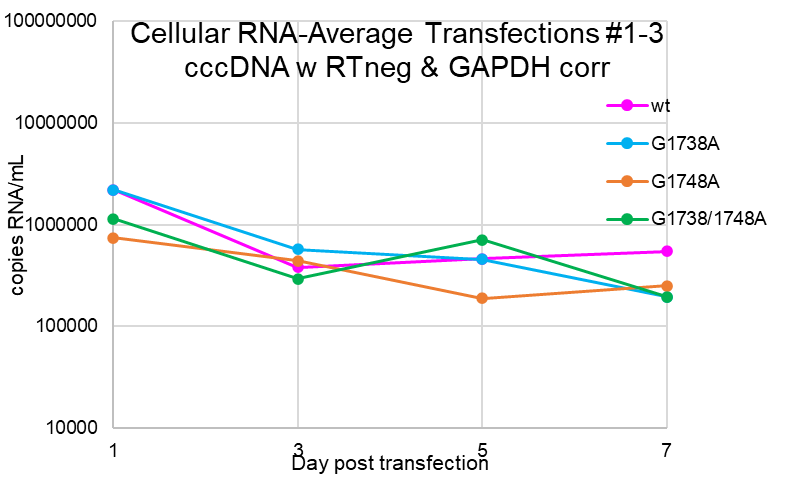

**Figure S6:** Average cellular RNA from 1.3mer HBV transfections of HepG2 cells. Cells harvested on days 1, 3, 5 and 7 and RNA extracted with Trizol, DNase digested. qPCR was performed using “cccDNA” primers (see Supplemental Table 4) to pick up all forms of HBV RNA. Data was corrected for qPCR results from both the reverse transcriptase negative samples and for the cell number as measured by the GAPDH internal control.

**SUPPLEMENTARY TABLES**

**Table S1:** Distribution of Peaks for CD data.

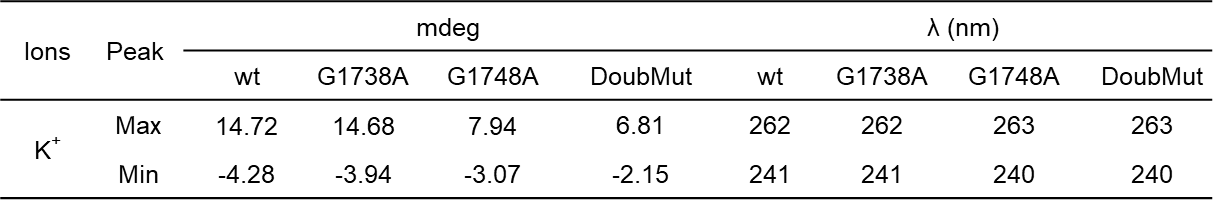

**Table S2:** Primers used for site-directed mutagenesis

| **Primer** | **Sequence (5′ – 3′)** |
| --- | --- |
| SDM-G1738A-For | GTTTAAGGACTGGGA**A**GAGTTGGGGGAGGAG |
| SDM-G1738A-Rev | CTCCTCCCCCAACTCT**T**CCCAGTCCTTAAAC |
| SDM-G1748A-For | CTGGGAGGAGTTGGGG**A**AGGAGATTAGGTTATTTG |
| SDM-G1748A-Rev | CAAATAACCTAATCTCCT**T**CCCCAACTCCTCCCAG |
| SDM-Doub-For | GTTTAAGGACTGGGA**A**GAGTTGGGG**A**AGGAG |
| SDM-Doub-Rev | CTCCT**T**CCCCAACTCT**T**CCCAGTCCTTAAAC |

**Table S3:** Primers used for direct, nested and quantitative PCRs

| **Primer** | **Sequence (5′ – 3′)** |
| --- | --- |
| cccDNA-D-For | ACTCCTGGACTCTCAGCAATG |
| cccDNA-D-Rev | GTATGGTGAGGTGAGCAATG |
| cccDNA-N-For /  HBV-totalRNA-F | AGGCTGTAGGCACAAATTGGT |
| cccDNA-N-Rev /  HBV-totalRNA-R | GCTTATACGGGTCAATGTCCA |
| TAKA-HBV-XD-F | GCATGGAGACCACCGTGAAC |
| TAKA-HBV-XD-R | GGAAAGAAGTCAGAAGGCAA |
| HBV-PreC/C-For2 | TCACCTCTGCCTAATCATC |
| HBV-PreC/C-Rev2 | GGAGTGCGAATCCACACTCC |
| GAPDH-For | ACCAACTGCTTAGCCC |
| GAPDH-Rev | CCACGACGGACACATT |
| Plasmid-Guo-For | AGGCTGTAGGCACAAATTGGT |
| Plasmid-Guo-Rev | GGTAGATCATACACATTGGC |
| HBV-PCP-For | GGTCTGCGCACCAGCACC |
| HBV-PCP-Rev | GGAAAGAAGTCAGAAGGCAA |
| HBV-core trans-For | TATCCCTTGGACTCACAAG |
| HBV-core trans-Rev | CAAGAATATGGTGACCCACA |

**Supplemental Table 4**: Summary of p-values for viral protein marker significance. All values are compared to the *wt* plasmid for corresponding days 1, 3, 5, and 7 post-transfection. p-values <0.05 are underlined.

| **Variant - Day** | **p-value** | | |
| --- | --- | --- | --- |
|  | **Supernatant HBsAg** | **Supernatant HBeAg** | **Cellular HBcAg** |
| G1738A-D1 | 0.0830 | 8.9E-05 | 0.06164 |
| G1738A-D3 | 0.0352 | 0.0028 | 0.0171 |
| G1738A-D5 | 0.0052 | 4.8E-06 | 0.0034 |
| G1738A-D7 | 0.0078 | 0.0003 | 0.0165 |
| G1748A-D1 | 0.2698 | 5.4E-07 | 0.0140 |
| G1748A-D3 | 0.0102 | 3.3E-06 | 0.0045 |
| G1748A-D5 | 0.0128 | 1.3E-07 | 0.0004 |
| G1748A-D7 | 0.1126 | 0.0004 | 0.0001 |
| G1738/1748A-D1 | 0.2831 | 6.0E-09 | 0.0007 |
| G1738/1748A-D3 | 0.0053 | 5.6E-06 | 1.9E-05 |
| G1738/1748A-D5 | 0.0066 | 0.1253 | 0.0003 |
| G1738/1748A-D7 | 0.0020 | 0.0010 | 2.3E-07 |

**SUPPLEMENTARY EXPERIMENTS**

**Transfection studies with linearized HBV**

A lab-derived wild type HBV plasmid was used that contained a complete copy of HBV genotype C virus inserted into a pUC19 vector, produced as previously described (56,57). The QuikChangeII site-directed mutagenesis kit (Agilent Technologies, Santa Clara, CA), was used to create the G1748A HBV mutant, using primers (University of Calgary DNA Synthesis Lab, Calgary, AB) described in the Supplementary data. The mutant HBV sequence was confirmed by Sanger sequencing. The original plasmid (*wt*) and the core-mutant (*G1748A*, “*mut*”) were amplified via an *E. coli* (Top10) (Thermo Fisher, Cat# C404010) amplification system and isolated using the GenElute Plasmid Miniprep kit (Sigma, Cat# PLN350). The purified plasmid was digested with BspQ1 restriction enzyme (New England Biolabs Ltd, Whitby, ON) and the 3.2 kb fragment isolated via gel electrophoresis to produce linear HBV genomes for transfection (HBV_wt-lin_ and HBV_mut-lin_). In addition to the above, an in-house green fluorescent protein (GFP)-pcDNA3.1(+) plasmid was used as control.

HepG2 cells were grown, plated and transfected with Lipofectamine3000 as described in the main manuscript text. On days 1, 3, 5 and 7, cells were imaged for GFP to determine % transfection. Immediately after imaging, the supernatant and cells were harvested for detection of HBV RNA, total HBV DNA, and/or viral antigens. Total RNA and DNA were extracted and viral antigens obtained from lysed cells per protocols described in the manuscript. Transfection experiments were performed in triplicate with duplicate samples for each condition per day, per experiment.

Detection and quantification of HBV RNA and DNA was performed using SYBR® Green fluorescence (BioRad, Cat# 1725124). An HBV-containing plasmid was used as a positive control and to produce the dilution series for the qPCR standards; water and “mock extracted” template and mock transfection were used as negative controls. A segment bridging the cut site of the linearized DNA (i.e., surrounding the HBV pre-core promoter region) was amplified using the primers, 5′-GGTCTGCGCACCAGCACC-3′ (HBV-PCP-For) and 5′-GGAAAGAAGTCAGAAGGCAA-3′ (HBV-PCP-Rev). This ensured detection of the original starting plasmid was not incorporated into the results. Samples were prepared in triplicate on the qPCR plate with concomitant standards and controls per run.

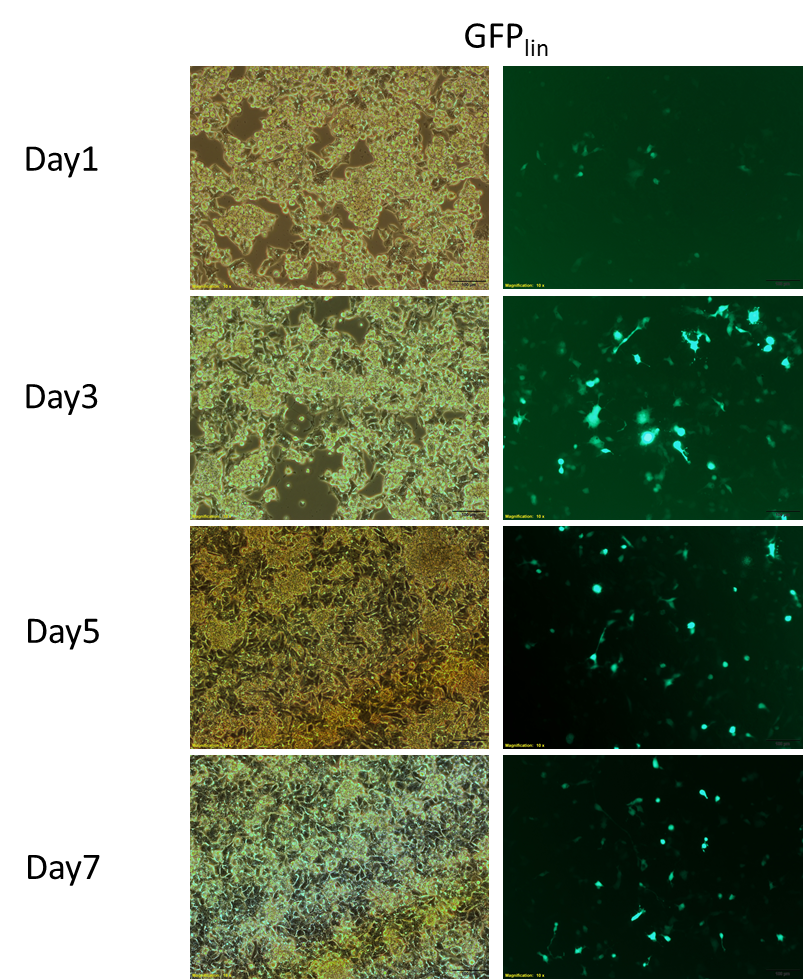

**Figure S7**: Confirmation of transfection efficacy of linearized plasmid. It is presumed that the linearized plasmid segment can enter with comparable efficiency as that of a circular plasmid and produce its subsequent product.

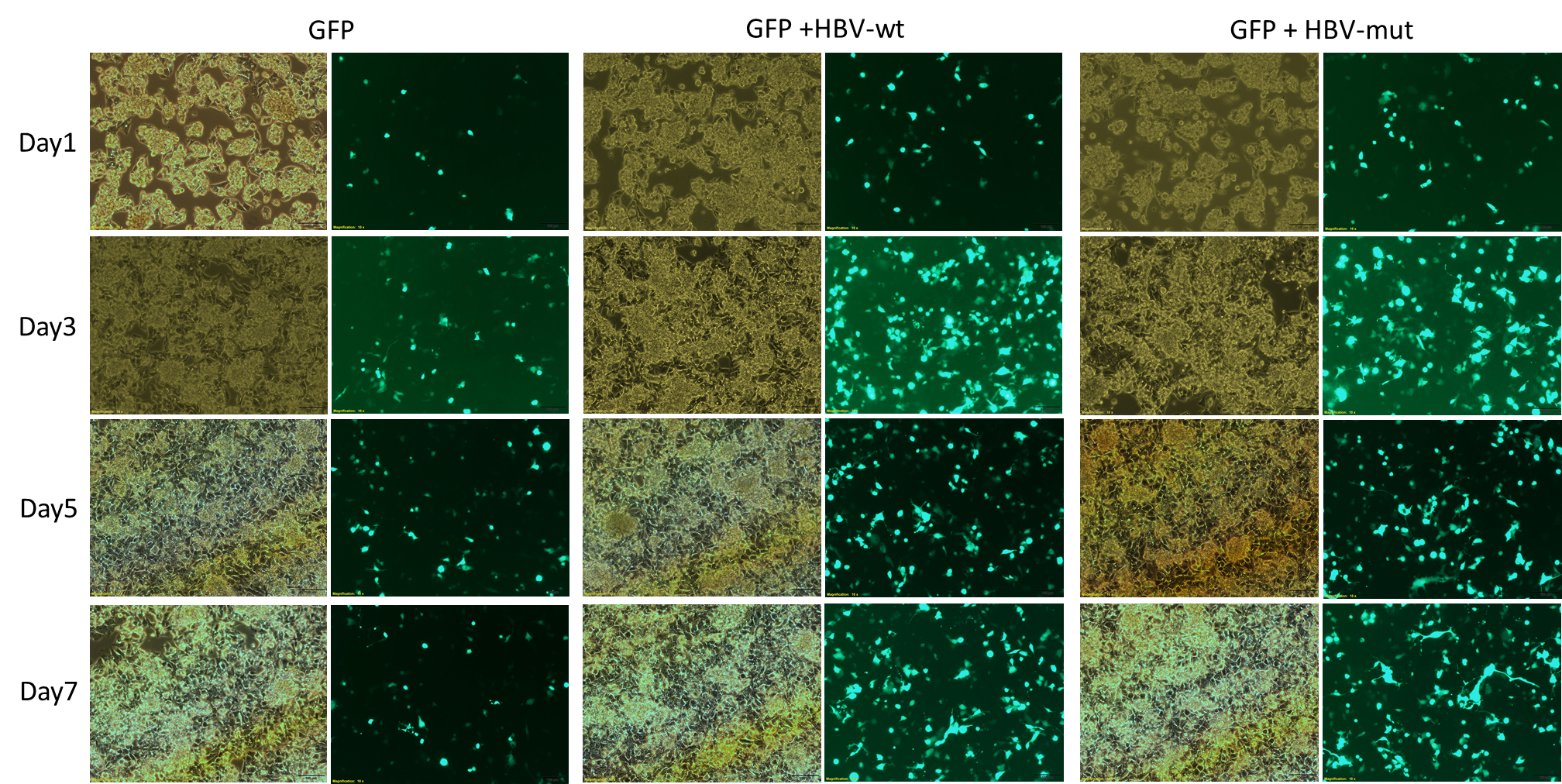

**Figure S8**: HepG2 transfection with linearized *wt* and *mut* (*G1748A)* HBV DNA. A separate GFP-based plasmid has been added to each well to monitor transfection efficiency in the cells, with the presumption that the linearized HBV would enter in a similar relative manner. Images were recorded on days 1, 3, 5, and 7 post-transfection.

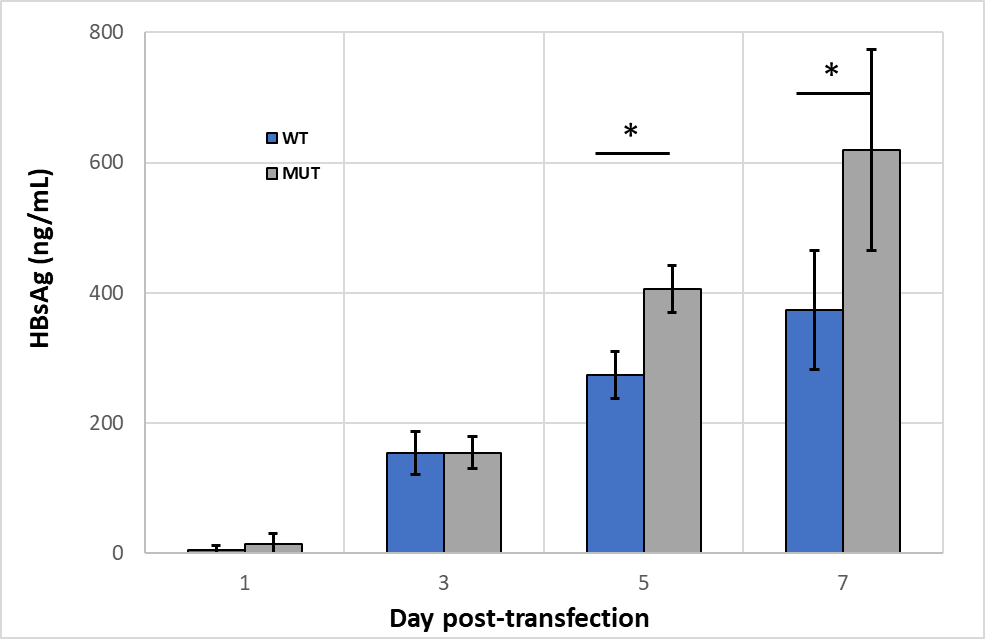

**Figure S9**: HepG2 transfection studies with *wt* and *mut* (*G1748A*) linearized HBV, harvested at days 1, 3, 5, and 7 post-transfection with sandwich ELISA for detection of supernatant HBsAg. p-values are 0.009 and 0.010, respectively, for days 5 and 7 post-transfection.

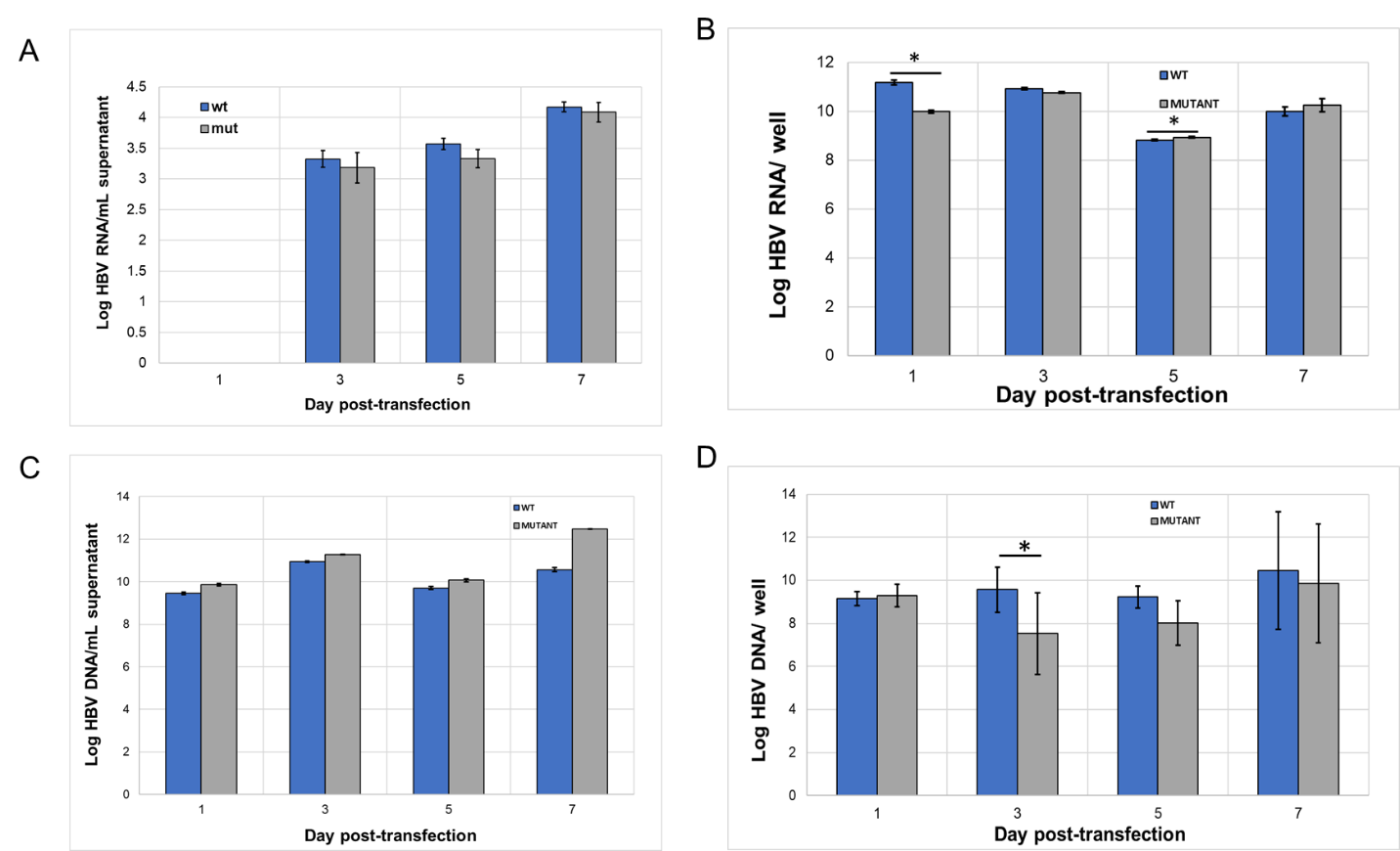

**Figure S10:** HepG2 transfection studies with *wt* and *mut* (*G1748A*) linearized HBV, harvested at days 1, 3, 5, and 7 post-transfection. Each plot is the averaged results of six replicates (duplicate samples per three transfections). Representative plots for total RNA in the supernatant **(A)** and cells **(B)**; total DNA in the supernatant **(C)** and cells **(D)** detected by qPCR. *p=<0.05.
